# Connectomics analysis reveals first, second, and third order thermosensory and hygrosensory neurons in the adult *Drosophila* brain

**DOI:** 10.1101/2020.01.20.912709

**Authors:** Elizabeth C. Marin, Ruairí J.V. Roberts, Laurin Büld, Maria Theiss, Markus W. Pleijzier, Tatevik Sarkissian, Willem J. Laursen, Robert Turnbull, Philipp Schlegel, Alexander S. Bates, Feng Li, Matthias Landgraf, Marta Costa, Davi D. Bock, Paul A. Garrity, Gregory S.X.E. Jefferis

**Affiliations:** Department of Zoology; University of Cambridge; Cambridge, CB2 3EJ; United Kingdom; Division of Neurobiology; MRC Laboratory of Molecular Biology; Cambridge, Cambridgeshire, CB2 0QH; United Kingdom; Department of Biology; Brandeis University; Waltham, MA 02454; USA; Janelia Research Campus; Howard Hughes Medical Institute; Ashburn, VA 20147; USA; Larner College of Medicine; University of Vermont; Burlington, VT 05405; USA

**Keywords:** Thermosensation, hygrosensation, connectomics, antennal lobe, projection neuron, lateral accessory calyx, circadian clock

## Abstract

Animals exhibit innate and learned preferences for temperature and humidity – conditions critical for their survival and reproduction. Here, we leveraged a whole adult brain electron microscopy volume to study the circuitry associated with antennal thermosensory and hygrosensory neurons, which target specific ventroposterior (VP) glomeruli in the *Drosophila melanogaster* antennal lobe. We have identified two new VP glomeruli, in addition to the five known ones, and the projection neurons (VP PNs) that relay VP information to higher brain centres, including the mushroom body and lateral horn, seats of learned and innate olfactory behaviours, respectively. Focussing on the mushroom body lateral accessory calyx (lACA), a known thermosensory neuropil, we present a comprehensive connectome by reconstructing neurons downstream of heating- and cooling-responsive VP PNs. We find that a few lACA-associated mushroom body intrinsic neurons (Kenyon cells) solely receive thermosensory inputs, while most receive additional olfactory and thermo- or hygrosensory PN inputs in the main calyx. Unexpectedly, we find several classes of lACA-associated neurons that form a local network with outputs to other brain neuropils, suggesting that the lACA serves as a general hub for thermosensory circuitry. For example, we find DN1 pacemaker neurons that link the lACA to the accessory medulla, likely mediating temperature-based entrainment of the circadian clock. Finally, we survey strongly connected downstream partners of VP PNs across the protocerebrum; these include a descending neuron that receives input mainly from dry-responsive VP PNs, meaning that just two synapses might separate hygrosensory inputs from motor neurons in the nerve cord. (249)

**HIGHLIGHTS:**

- Two novel thermo/hygrosensory glomeruli in the fly antennal lobe
- First complete set of thermosensory and hygrosensory projection neurons
- First connectome for a thermosensory centre, the lateral accessory calyx
- Novel third order neurons, including a link to the circadian clock

## INTRODUCTION

Temperature and humidity are two interrelated environmental variables with dramatic effects on animal physiology and survival. Animals exhibit intrinsic species-specific preferences for the temperature and humidity levels compatible with their survival and reproduction [1]. However, these preferences can be modulated by state and experience; vinegar flies prefer higher levels of relative humidity when dehydrated [2], and nematodes starved at low humidity subsequently prefer higher humidities, and vice versa [3]. Temperature and/or humidity also cue essential behavioural programmes; for example, female mosquitoes use heat and humidity to locate warm-blooded hosts for blood feeding [4, 5], and temperature entrains the circadian clock in organisms from molds to mammals to drive daily rhythms of physiology and behaviour [6, 7].

Whilst fundamental for survival, the neuronal mechanisms of thermosensation and hygrosensation are not well understood. In mammals, the molecules and circuits underlying body temperature homeostasis are still active areas of investigation despite significant progress on thermal nociception [8]. Small ectotherms, such as insects, are especially vulnerable to temperature and humidity extremes. Decades of study have identified peripheral thermo- and hygrosensors and the brain regions they innervate, but how sensory inputs are processed to drive behaviour is largely unknown [9]. Thermo- and hygrosensory neurons have been found on the antennae of diverse insect species, with individual sensilla commonly containing one cold air-responsive, one dry air-responsive, and one moist air-responsive neuron [10]. These neurons (like olfactory sensory neurons) project via the antennal nerve to the antennal lobe in the brain, where they innervate stereotyped subcompartments called glomeruli [11]. Projection neurons (PNs) relay information from these glomeruli to higher brain regions including the mushroom body calyx (CA) and lateral horn (LH) of the protocerebrum [12], areas involved in learned and innate olfactory behaviours, respectively [13].

Recent studies in *Drosophila melanogaster* have begun to provide detailed insights into the molecular mechanisms and higher order neural circuits underlying insect thermosensation and hygrosensation. In flies, the third antennal segment (funiculus) features two structures housing thermo- and hygrosensory neurons: the arista and the sacculus [14, 15]. The arista is a feathery protrusion containing three (sometimes four) pairs of phasic thermosensory neurons, one transiently activated by warming and the other by cooling [16, 17]. The sacculus is a multi-chambered invagination [18] containing several types of sensilla, some of which resemble thermosensory or hygrosensory sensilla. In particular, Chamber I contains 5 - 7 poreless basiconic sensilla, each housing two or three sensory neurons, and Chamber II contains 9 - 10 poreless coeloconic sensilla, each housing three sensory neurons [19]. Consistent with sensilla morphology, dry-responsive neurons have been identified in chambers I and II [20, 21] and humid-responsive neurons in Chamber II [22, 23].

At the molecular level, in the arista, heating-activated neurons detect temperature via the gustatory receptor isoform Gr28b.d [24] and cooling-activated neurons via the ionotropic receptor Ir21a, which requires co-receptors Ir25a and Ir93a [17]. Ionotropic receptors are also expressed in the sacculus, with Ir40a mediating dry sensation [20–22] and Ir68a humid sensation [22, 23], in both cases with Ir25a and Ir93a [20–22], suggesting widespread dependence on ionotropic receptor signalling.

At the circuit level, the funicular thermosensory and hygrosensory neurons primarily innervate ventroposterior (VP) antennal lobe glomeruli. Aristal thermosensory neuron axons arborise unilaterally in either VP2 (warming) or VP3 (cooling) [16, 25]. The saccular Ir40a-expressing, dry air-responsive neurons innervate VP4 (the “arm”) [20,21,26], in both antennal lobes, and Ir68a-expressing, humid air-responsive neurons project ipsilaterally to VP5 [22, 23]. In addition, putative thermosensors in Chamber II, which also express Ir40a, project to VP1 (the “column”) [20,21,26] (Fig 1A). VP1 neurons have been variously reported to respond weakly to cooling [20], dry air [21], or ammonia [27] and may represent evaporative cooling.

**Figure 1:**
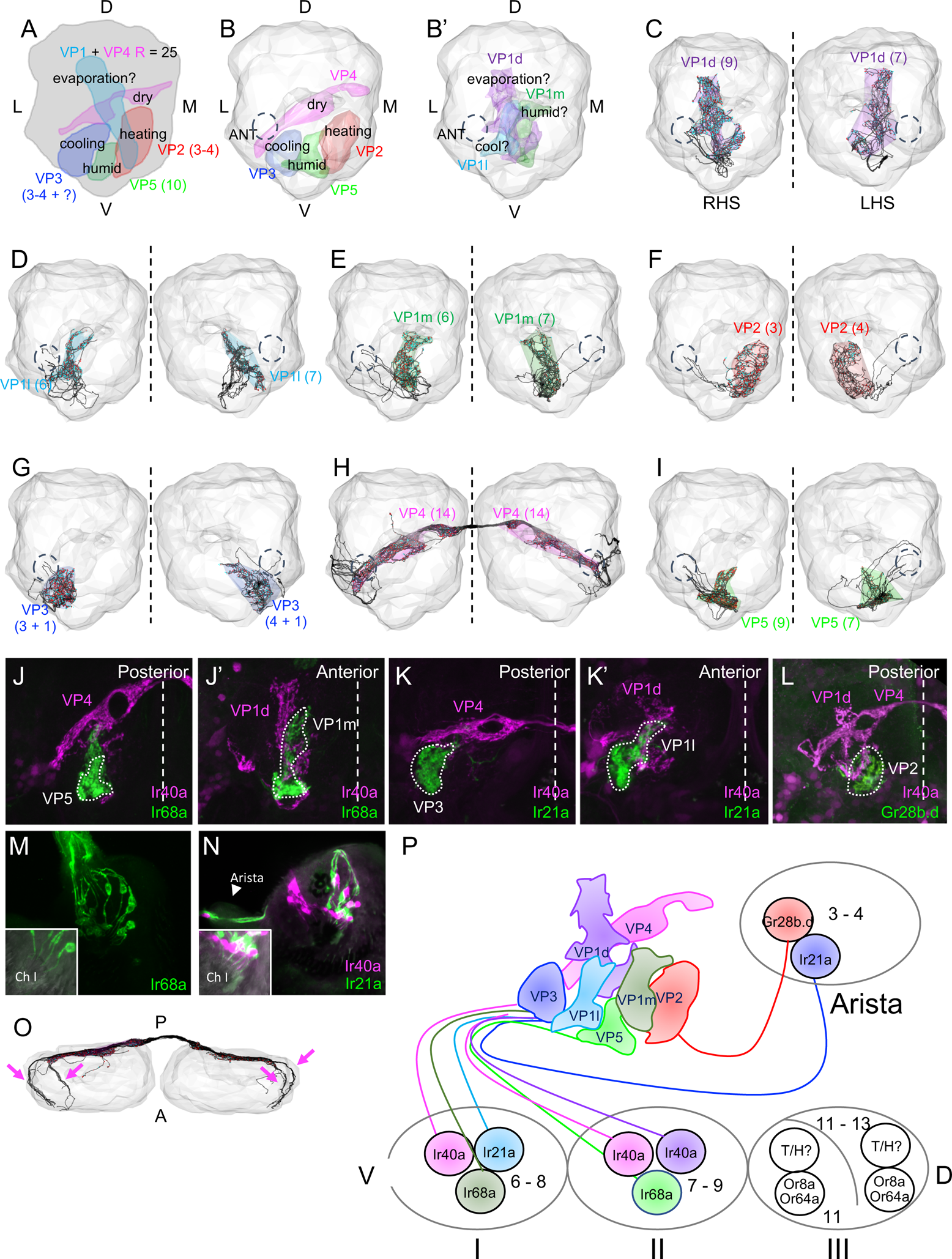
Sensory neurons define seven distinct glomeruli in the ventroposterior antennal lobe. **A.** Frontal view illustration of the VP antennal lobe summarising the sensory modalities, locations, and numbers of the five thermosensory and hygrosensory neuron classes reported in previous studies. D: Dorsal. L: Lateral. **B - B’.** Glomerular meshes generated from sensory neurons reconstructed in the right hemisphere of FAFB. Dashed circle labelled ANT: right antennal nerve. **B.** Posterior meshes: VP2, VP3, VP4, and VP5. **B’.** Anterior meshes: VP1d, VP1l, and VP1m. **C – I.** Antennal lobes with glomerular meshes enclosing FAFB reconstructions of individual VP sensory neurons (in black, with presynaptic sites in red and postsynaptic sites in cyan, and number of neurons indicated in parentheses; right hemisphere shown on left; dashed circles: antennal nerves; dashed line: the midline). **C.** VP1d. **D.** VP1l. **E.** VP1m. **F.** VP2. **G.** VP3 (3 - 4 aristal and one additional). **H.** VP4. **I.** VP5. **J – L.** Frontal views of receptor expression data in the antennal lobe. White dashed line: the midline. **J – J’.** VP1m and VP5 sensory neurons express Ir68a (green) but not Ir40a (magenta). **K – K’.** VP1l and VP3 sensory neurons express Ir21a (green) but not Ir40a (magenta). **L.** VP2 sensory neurons express Gr28b.d (green), but VP1l sensory neurons do not. Magenta: Ir40a. **M.** Both Chamber I (Ch I) and Chamber II sensory neurons express Ir68a (green) (16.14 + 1.77, n = 7). Insert: bright field overlay with GFP. **N.** Chamber I (Ch I) sensory neurons express Ir21a (green) but not Ir40a (magenta) (7.18 + 1.25, n = 11). Insert: bright field overlay with GFP. Arrowhead: aristal Ir21a+ neurons (to VP3). **O.** Dorsal view of VP4 HRNs. Arrows: two distinct axon bundles from each antennal nerve. **P.** A model of the organisation of thermosensory and hygrosensory neurons in the three-chambered sacculus and arista, with the specific receptors they express and the VP antennal lobe glomeruli they innervate.

Several classes of PNs have been reported to relay information from VP glomeruli to higher brain centres in *Drosophila*. A slow-adapting, cool-sensing VP3 PN takes the mediolateral antennal lobe tract (mlALT) to the lateral accessory calyx (lACA) of the mushroom body [28, 29]. Fast-adapting thermosensory PNs take the lateral antennal lobe tract (lALT) to target the posterior lateral protocerebrum (PLP) and posterior slope (PS) [29, 30]. One PN innervating several VP glomeruli takes the medial antennal lobe tract (mALT) to the PLP [23]. These studies indicate that thermo- and hygrosensory pathways project to multiple brain regions and suggest that these inputs are integrated at the level of the antennal lobe PNs.

Despite these advances, our current description of thermo- and hygrosensory systems in *Drosophila* is incomplete. The sacculus contains multiple sets of uncharacterised sensory neurons which likely innervate as-yet unknown antennal lobe glomeruli. Moreover, uniglomerular PNs have not yet been reported for several of the known VP glomeruli, suggesting more remain to be identified. Finally, little is known about higher order neurons, apart from Kenyon cells (mushroom body intrinsic neurons) of the α’/β’ subtype being likely targets of the slow-cool VP3 PN [31].

To obtain a more comprehensive view of the thermo- and hygrosensory system, we took advantage of FAFB, a Full Adult Fly Brain electron microscopy (EM) volume that permits whole neuron reconstruction and synapse annotation [32]. We identified the five previously reported VP glomeruli and two novel glomeruli with their respective sensory neurons, showing they are thermo- or hygrosensory. We also reconstructed every PN projecting from VP glomeruli to higher brain centres, finding them to be highly diverse in tract, neuropil target, and neuroblast of origin. We elucidated the “connectome” of the lACA, finding that this structure not only provides input to both dedicated and integrative Kenyon cells but also serves as a hub for the processing of thermosensory information and a link to the circadian clock. Finally, we identified additional novel third order neurons, one of which is a descending neuron that relays hygrosensory information to the ventral nerve cord. Together, these data provide a comprehensive first- and second-order layer analysis of *Drosophila* thermo- and hygrosensory systems, and an initial survey of third order neurons which subsequently modulate behaviour.

## RESULTS

### Sensory neurons define two novel glomeruli in the ventroposterior antennal lobe

Five glomeruli in the ventroposterior (VP) antennal lobe that receive thermosensory or hygrosensory inputs from the antennae have been previously described anatomically and/or functionally (Fig. 1A). To define these glomeruli in FAFB, we reconstructed putative sensory neurons in both hemispheres.

We first identified two thermosensory glomeruli, VP2 (heating) and VP3 (cooling), in the posteriormost AL (Fig. 1B). Three unilateral thermosensory neurons (TRNs) with complex arbours on the right-hand side (RHS) and four on the left-hand side (LHS) innervated a medial glomerulus, VP2 (Fig. 1F). Four unilateral sensory neurons on the RHS and five on the LHS innervated a lateral glomerulus, VP3 (Fig. 1G), with all but one forming complex arbours (Fig. S1E’ - E’’) and the last traveling separately in the antennal nerve and forming a simpler arbour (Fig. S1E’’’). Morphological clustering with NBLAST [33] of unilateral VP sensory neurons paired the simpler VP3 TRNs on each side with one another, within the larger VP3 TRN group (Fig. S2A), implying that they are of the same type and distinct from the complex VP3 TRNs. Moreover, the simpler VP3 TRN on the RHS provided most of the input to the slow-cool VP3 PN projecting to the lACA, whereas the others provided input to a distinct class of VP3 PNs (Fig. S2B). These results are consistent with reports of three or four pairs of heating-responsive and cooling-responsive aristal sensory neurons that project to VP2 and VP3, respectively [16, 17], and primarily non-aristal cold input to the slow-cool VP3 PN [29].

We next identified two known hygrosensory glomeruli, VP4 (dry air) and VP5 (humid air), in the RHS AL (Fig. 1B). We traced 28 bilateral hygrosensory neurons (HRNs), 14 from each antennal nerve, that collectively defined the VP4 glomerulus (Fig. 1H). These VP4 HRNs travelled in two distinct areas of each antennal nerve (Fig. 1O), perhaps reflecting their origins in sacculus chambers I vs. II. Between VP2 and VP3, we found 9 unilateral sensory neurons on the RHS and 7 on the LHS occupying VP5 (Fig. 1I), consistent with previous reports [22]. Since VP4 and VP5 HRNs are found in the same triad sensilla in Chamber II, this suggested that 9 of the 14 VP4 HRNs on the RHS and 7 on the left originated in Chamber II, with the remainder coming from Chamber I.

To our surprise, the axons of unilateral sensory neurons in the vicinity of the previously described glomerulus VP1 formed not one, but rather three distinct but adjacent populations (Fig. 1B’) which we designated according to their relative positions: VP1d (dorsal, Fig. 1C), VP1l (lateral, Fig. 1D), and VP1m (medial, Fig. 1E). Morphological clustering confirmed that the unilateral VP sensory neurons constituted six distinct classes: VP1d, VP1l, VP1m, VP2, VP3, and VP5 (Fig. S2A). Sensory neurons from VP1d, VP1l, and VP1m each formed synaptic connections with other neurons within the same class, but not with neurons from either of the other two classes, consistent with glomerular compartmentalisation (Fig. S2C).

Light-level studies (relying on IR40a-specific antibodies and drivers) had identified a total of 25 saccular sensory neurons projecting to VP1 and VP4 combined [21,22,26]. VP1d HRNs most closely resembled the Ir40a-positive “VP1” neurons in morphology and location, and the total number of VP1d and VP4 HRNs on the RHS of FAFB was 23 and on the LHS 21. Also, we reconstructed the same numbers of VP1d and VP5 neurons in a given antennal lobe (Fig. 1C, I), suggesting that these classes might cohabit the triad sensilla in Chamber II, along with the VP4 HRNs [19,22,23].

We hypothesised that the novel VP1l and VP1m glomeruli expressed distinct receptors from VP1d and likely represented distinct sensory modalities. To investigate this possibility, we generated novel tools to label neurons that express other receptors previously associated with thermo- and hygrosensation (see Methods). While Ir68a expression had been reported in humid air-responsive VP5 HRNs, using these new tools we also observed expression in VP1m (Fig. 1J-J’). We also observed expression of Ir21a (Fig. 1K-K’) but not Gr28b.d (Fig. 1L) in VP1l neurons, suggesting that VP1l neurons are cool- or cooling-responsive. These expression data, together with their characteristic neuronal and sensillar morphologies, are consistent with these new VP sensory neurons being thermo- or hygrosensory in nature (Fig. 2B’).

**Figure 2:**
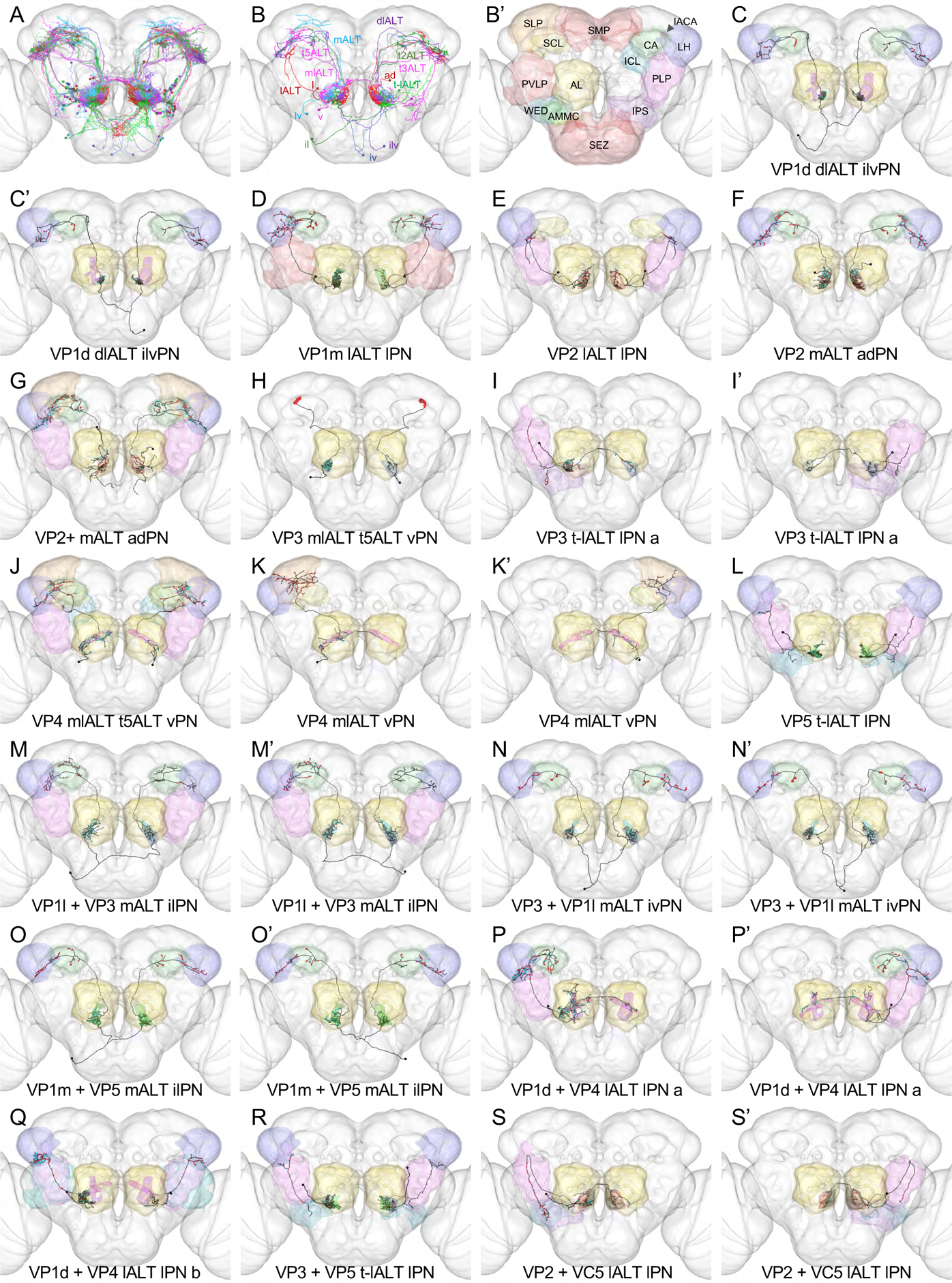
Diverse unique ventroposterior projection neurons relay information from the antennal lobe to numerous higher brain centres. **A.** Frontal view of all VP PNs, colour-coded by primary glomerulus. **B.** Cartoon summarising the tracts and neuroblast lineages of VP PNs. **B’.** Cartoon summarising the target brain neuropils of VP PNs, colour-coded to correspond with all other figure panels. AL: antennal lobe. AMMC: antennal mechanosensory and motor centre. CA: mushroom body calyx. ICL: inferior clamp. IPS: inferior posterior slope. lACA: lateral accessory calyx. LH: lateral horn. PLP: posterior lateral protocerebrum. PVLP: posterior ventral protocerebrum. SCL: superior clamp. SEZ: suboesophageal zone. SLP: superior lateral protocerebrum. SMP: superior medial protocerebrum. WED: wedge. **C – S’.** Frontal views of individual VP PNs, with glomeruli colour-coded as in Fig. 1B - B’ (non-VP glomeruli in grey). For more details re: individual neurons, please see Table 1.

We observed the same numbers of VP1l and VP1m neurons for a given antennal lobe (Fig. 1D, E), suggesting common sensillar origins. Consistent with this model, Ir68a (Fig. 1M) and Ir21a expression (Fig. 1N) were each detected in ∼7 neurons of Chamber I. These neurons would form a triad with the Ir40a-expressing neurons in Chamber I (Fig. 1N) projecting to VP4.

In summary, our data along with published expression data support a model in which VP2 and VP3 are innervated by heating-responsive and cooling-responsive sensory neurons from the aristae, while five additional VP glomeruli are innervated by sensory neurons from the sacculus (Fig. 1P). Sensory neurons from 7 - 9 triad sensilla in Chamber II innervate VP4 (dry), VP5 (moist), and VP1d (putative evaporative cooling). Sensory neurons from 6 - 8 sensilla in Chamber I innervate VP4 (dry), VP1l (cool/cooling), and VP1m (possibly humid).

### A diverse population of projection neurons relays activity from the ventroposterior antennal lobe to higher brain centres

Our next goal was to identify the projection neurons (PNs) that relay activity from the ventroposterior antennal lobe glomeruli to higher brain centres. We reconstructed a total of 76 PNs that received input primarily from one or more of the seven VP glomeruli, henceforth referred to as VP PNs (Table 1, Fig. 2A). We completely reconstructed the axons of 39 VP PNs on the RHS, 19 of which were novel, and reconstructed 37 VP PNs on the left to identification (Fig. 2C - U, Fig. 3, Fig. S3A-C). We consistently found the same 32 classes of VP PNs in both hemispheres (based on glomerulus, tract, and lineage and confirmed by clustering with NBLAST), strongly suggesting that they were morphologically stereotyped (Fig. S3B).

**Table 1:** V**e**ntroposterior **projection neurons.** NAME: our designation, based on glomerulus innervated, antennal lobe tract, and neuroblast lineage. PANEL: corresponding panel in this paper. SKID_R: right-hand side skeleton ID. SKID_L: left-hand side skeleton ID. VFB_ID: VFB identification number. FC_MATCH: corresponding single-cell clone in FlyCircuit, based on NBLAST. PREV_NAMES: names previously assigned to this neuron in the literature. DRIVERS: sparse driver line(s) previously reported to label this neuron. NT: likely neurotransmitter, based on neuroblast lineage and tract. OTHER: Previous publications featuring reconstructions of this neuron in FAFB.

| NAME | PANEL | SKID_R | SKID_L | VFB_ID | FC_MATCH | PREV_NAMES | DRIVERS | NT | OTHER |
| --- | --- | --- | --- | --- | --- | --- | --- | --- | --- |
| VP1d dIALT iIvPN | 2C, C' | 192423 | 203504 |  | VGlut-F-000344 |  |  |  | [32] |
| VP1m IALT IPN | 2D | 11524119 | 3078 |  | Cha-F-500056 | unPN-2 [37]; IPN1 [34] |  | GABA? |  |
| VP2 IALT IPN | 2E | 5672990 | 13154015 |  |  | unPN-4 [37] | VT046265 | GABA? |  |
| VP2 mALT adPN | 2F | 57516 | 9159128 |  | Gad1-F-000090 | VP2 uACT1 [25] |  | ACh | [32] |
| VP2+ mALT adPN | 2G | 1712057 | 8864990 |  |  |  |  | ACh |  |
| VP3 mIALT t5ALT vPN | 2H | 5471515 | 561100 |  | Fru-M-100050 | AL-t5PN1[28]; slow-cool PN [29] | R60H12 [30] | GABA |  |
| VP3 t-IALT IPN a | 2I, I' | 4869882 | 13159741 |  |  | fast-cool PN [29] | R95C02 [29] | GABA? |  |
| VP4 mIALT t5ALT vPN | 2J | 1149173 | 9039579 |  | VGlut-F-300564 |  |  | GABA | [32] |
| VP4 mIALT vPN | 2K, K' | 2484510 | 12805737 |  |  |  |  | GABA |  |
| VP5 t-IALT IPN | 2L | 4672650 | 13053419 |  |  |  | R95C02 | GABA? |  |
| VP1I + VP3 mALT iIPN | 2M, M' | 57495 | 57487 |  | Cha-F-100277 | VP3 bilateral PN [25] |  | ACh |  |
| VP3 + VP1I mALT iIvPN | 2N, N' | 45882 | 37513 |  |  |  | VT26020 [30] | ACh | [32] |
| VP1m + VP5 mALT iIPN | 2O, O' | 57503 | 46105 |  | Gad1-F-300444 | R24G07 hPN [23] | R24G07 [23] | ACh | [32] |
| VP1d + VP4 IALT IPN a | 2P, P' | 23005 | 9716626 |  |  | biPN-2 [37] |  | GABA? | [32] |
| VP1d + VP4 IALT IPN b | 2Q | 3908507 | 9740174 |  | fru-M-000094 | VP3 oACT [25] |  | GABA? |  |
| VP3 + VP5 t-IALT IPN | 2R | 4671556 | 9704839 |  | Gad1-F-400223 | unPN-6 [37]; cool IALT PN [30] | R95C02 [30] | GABA? |  |
| VP2 + VC5 IALT IPN | 2S, S' | 2600341 | 10515783 |  | Cha-F-100163 | biPN-3 [37] |  | GABA? |  |
| VP3 t-IALT IPN b#1 | 3A | 11234968 | 13159728 |  |  |  |  | GABA? |  |
| VP3 t-IALT IPN b#2 | 3A' |  | 13159715 |  |  |  | R95C02 | GABA? |  |
| VP5 + VP2 t-IALT IPN #1 | 3B | 5946848 | 13159719 |  | Cha-F-100123 | warm-PN [29] | R95C02 [29] | GABA? |  |
| VP5 + VP2 t-IALT IPN #2 | 3B' | 4876532 |  |  | Cha-F-100123 | warm-PN [29] | R95C02 [29] | GABA? |  |
| VP t-IALT IPN #1 | 3C, C' | 7017522 | 5946637 |  |  |  |  | GABA? |  |
| VP t-IALT IPN #2, 3 | 3C'', C''' | 1356477 | 5582907 |  |  |  |  | GABA? |  |
| VP1I+ mALT iIvPN #1 | 3D | 57200 | 13492671 |  |  |  |  | ACh |  |
| VP1I+ mALT iIvPN #2 | 3D' | 56999 | 7463926 |  |  |  |  | ACh |  |
| VP1I+ mALT iIvPN #3 | 3D'' | 57224 | 4177248 |  |  |  |  | ACh |  |
| VP5+ SEZ adPN | 3E | 14003783 | 9511430 |  |  | poly[emb] adPN [38] |  | ACh | [32] |
| VP4 + VP2 + VL1 t3ALT IPN | 3F, F' | 1664544 | 4631122 |  |  | warm-cool PN [29] | R54A03 [29] | GABA? |  |
| VP1m+ t2ALT lvPN | 3G | 192430 | 9590169 |  |  |  |  |  |  |
| VP1m + VP2 t2ALT lvPN | 3G' | 4058666 | 9588978 |  |  |  |  |  |  |
| VP1m + VP2 mALT lvPN a#1 | 3H | 57122 | 9101937 |  |  |  |  | ACh |  |
| VP1m + VP2 mALT lvPN a#2 | 3H' | 57059 | 12681995 |  |  |  |  | ACh |  |
| VP1m + VP2 mALT lvPN a#3 | 3H'' | 57114 | 13931917 |  |  |  |  | ACh |  |
| VP1m + VP2 mALT lvPN b | 3I | 57063 | 4495405 |  |  |  |  | ACh |  |
| VP1m + VP2 mALT lvPN c | 3J | 57138 | 11052217 |  |  |  |  | ACh |  |
| VP mALT lvPN | 3K | 57142 |  |  |  |  |  | ACh |  |
| VC5+ t3ALT IPN#1 | 3L | 1363077 | 5122761 |  | VGlut-F-700270 | slow hot and cold t3ALT PN [30] | R84E08 [30] | GABA? |  |
| VC5+ t3ALT IPN#2 | 3L' | 8825246 | 13950267 |  | VGlut-F-100102 | slow hot and cold t3ALT PN [30] | R84E08 [30] | GABA? |  |
| VP2 SEZ mALT lvPN #1 | 3M, M' | 57166 | 7423431 |  |  |  |  | ACh |  |
| VP2 SEZ mALT lvPN #2 | 3M'' | 57146 |  |  |  |  |  | ACh |  |
| VC5+ t-IALT IPN | 3N | 8406430 |  |  |  |  |  | GABA? |  |
| VP1d+ mIALT vPN#1 | S3A | 3813438 | 5643689 |  |  |  |  | GABA? |  |
| VP1d+ mIALT vPN#2 | S3A' | 3813424 | 13272972 |  |  |  |  | GABA? |  |

**Figure 3:**
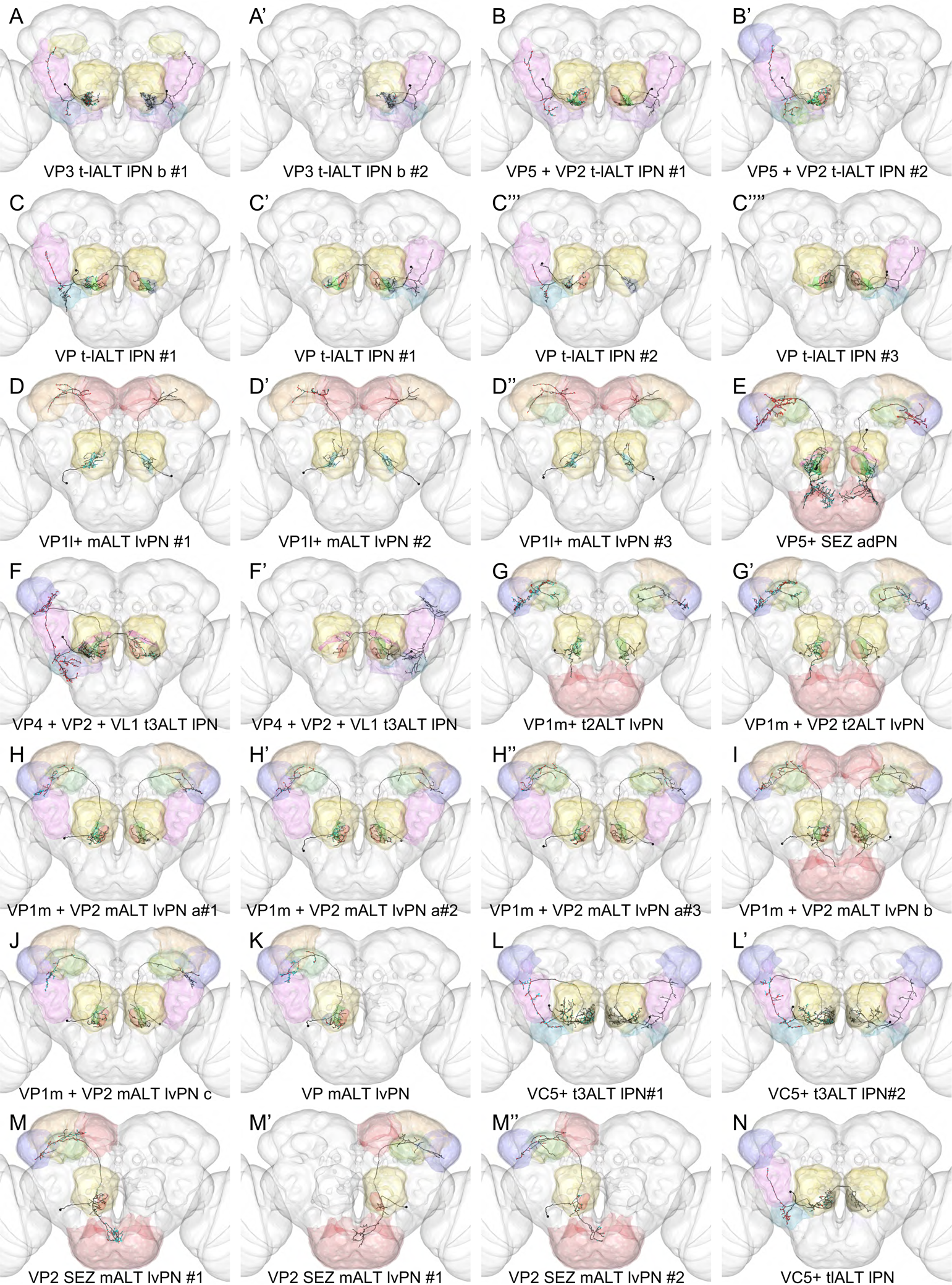
A diverse population of ventroposterior projection neurons relays information from the antennal lobe to numerous higher brain centres. A - Y. Frontal views of individual VP PNs, with glomeruli colour-coded as in Fig. 1B - B’ (non-VP glomeruli in grey) and brain neuropils colour-coded as in Fig. 2B’. For more details re: individual neurons, please see Table 1.

Our reconstructed VP PNs constituted a highly diverse population, projecting to higher brain centres via any of seven antennal lobe tracts: the mALT, mlALT, lALT, a small posterior tract leaving the lALT that we dubbed the transversal-lALT (t-lALT), t2ALT, t3ALT, or t5ALT (Fig. 2B, Table 1) [34]. Like canonical excitatory olfactory PNs, many VP PNs targeted the CA and LH, where their terminals were largely confined to specific domains (Fig. 4A - A’). However, VP PNs also targeted additional neuropils, particularly the SLP, PLP, wedge, and the lACA (Fig. 2B’).

**Figure 4:**
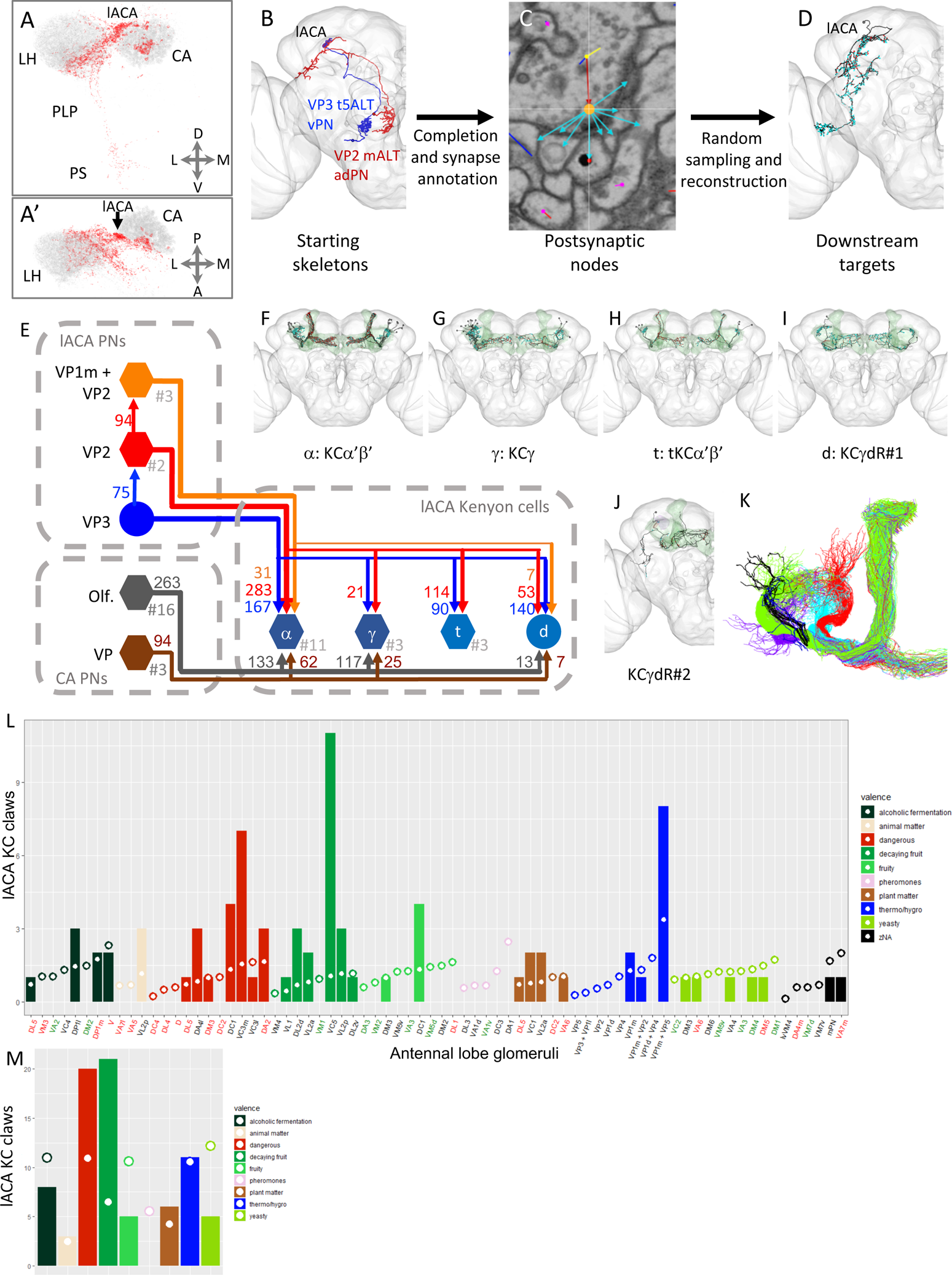
Both dedicated and mixed modality Kenyon cells receive thermosensory input from the lateral accessory calyx. A – A’. Frontal (A) and dorsal (A’) views of VP PN presynapses (red) vs. olfactory PN presynapses (grey). CA: mushroom body main calyx. LH: lateral horn. PLP: posterior lateral protocerebrum. PS: posterior slope. **B – D.** Workflow for randomised downstream sampling from key VP PNs in the lACA. **B.** Frontal view of reconstructed VP PNs used for sampling from the lACA in the right hemisphere of FAFB. **C.** Annotation of presynapses and associated postsynapses from these two VP PNs in the lACA. Orange: presynapse. Cyan arrow: edge to associated postsynapse. **D.** Frontal view of an example target neuron reconstructed during the sampling process. **E.** Partial circuit diagram illustrating connectivity between lACA-associated Kenyon cells receiving at least 5 inputs in the lACA and their VP PN and olfactory inputs. Circular nodes represent individual neurons, and hexagonal nodes represent pooled neurons, with the number pooled after the hashtag. Synapse numbers correlate with line thicknesses. lACA PNs: PNs targeting the lACA. CA PNs: Olfactory and VP PNs targeting KC dendrites in the main calyx. **F – J.** Frontal views of lACA-associated Kenyon cell reconstructions. For more details re: individual neurons, please see Fig. S5. Green: whole mushroom body. Red: lACA. Purple: dACA (**J** only). **K.** lACA-associated KCs (black) derive from one of four mushroom body neuroblast lineages (coloured). **L.** Expected (circles) vs. observed (bars) numbers of lACA KC dendritic claws receiving input from olfactory and VP PN partners in the CA (colour-coded by valence, key on right). Red labels: aversive. Green labels: attractive. Black labels: neutral or unknown. **M.** Summary of expected (circles) vs. observed (bars) numbers of lACA KC dendritic claws receiving input of various valences (colour-coded by valence, key on right) from olfactory and VP PN partners in the CA.

Based on their primary soma tracts, the VP PNs belonged to seven distinct neuroblast lineages: AL anterodorsal (ad), lateral (l), ventral (v), or lateroventral (lv) [35], or the previously unidentified inferior lateral (il), inferior ventral (iv), or inferior lateroventral (ilv) [36]. The lateral lineage generated the most VP PNs [37] and exhibited the most diversity in axon tract (Table 1). Notably, a subset of these lPNs relayed information from VP5, VP2, and/or VP3 to the PLP, PS, and/or wedge via the transversal-lALT (Figs. 2I-I’, 2L, 3A-C’’’’) and likely mediate rapid responses to changes in humidity and/or temperature [29].

Six of the seven VP glomeruli were innervated by at least one unique, uniglomerular VP PN (Figs. 2C-F, 2H-L). This validates the boundaries of the VP glomeruli and particularly our distinction between VP1d, VP1l, and VP1m. However, we did not find any bona fide uniglomerular VP1l PNs, only several multiglomerular lvPNs that primarily innervated VP1l and thus were designated VP1l+ (Fig. 3D-D’’). Also, only a portion of the VP1d glomerulus was innervated by its uPN (Fig. 2C-C’), suggesting the potential for localised contacts.

Unexpectedly, several unique VP PNs innervated two or more distinct glomeruli. Three were bilaterally symmetric, innervating VP3 and VP1l (Fig. 2M-N’) or VP1m and VP5 (Fig. 2O - O’), while two asymmetric lPNs innervated VP1d and VP4 (Fig. 2P-Q). These could integrate the activity of two types of cooling-, moist-, or dry/evaporation-responsive sensory neurons, respectively. We reconstructed one unique lPN innervating VP2 and VC5 (Fig. 2S-S’), and another innervating VP2, VP4, and VL1 (Fig. 3F-F’) reportedly activated by both warming and cooling air [29]. We also identified a multiglomerular VP5+ adPN [38] with extensive innervation of the SEZ (Fig. 3E), and three RHS lvPNs innervating the VP2 and the SEZ (Fig. 3M-M’’), presumably integrating humidity or temperature with taste.

We reconstructed at least two distinct classes of VP1m + VP2 PNs: a pair taking the t2ALT to the CA, LH, and SLP (Fig. 3G-G’) and a group of lvPNs taking the mALT to the CA and LH (Fig. 3H-J). We also found a pair of multiglomerular VC5+ t3ALT lPNs (Fig. 3K-K’) that had been previously implicated in thermosensation [30] and a VC5+ lPN traveling near VP PNs in the t-lALT (Fig. 3N). Finally, we noted that numerous vPNs innervated VP1d along with other VP or olfactory glomeruli (Fig. S3A-A’). We speculate that these common combinations of glomeruli could represent stimulus combinations of particular behavioural importance.

In summary, we reconstructed 32 types of VP PNs, only 9 of them uniglomerular. We identified single-cell clones in <u>FlyCircuit</u> for 11 types (Table 1, FC_MATCH) and sparse GAL4 or LexA driver lines in <u>Fly Light</u> for 10 (Table 1, DRIVERS). We found anatomical (and for a subset, functional) descriptions of 14 types in the *Drosophila* neurobiology literature (Table 1, PREV_NAMES). However, elucidation of our 19 novel VP PN types awaits future physiological and behavioural studies.

### VP PNs relay thermosensory information to Kenyon cells innervating the lateral accessory calyx

How is thermosensory and hygrosensory information organised in higher brain centres, and what are the implications for multimodal sensory integration and behaviour? Although the VP PNs target many different neuropils, the majority project to the mushroom body calyx and/or LH. Moreover, most VP PN terminals are restricted to the anterior of the calyx, either in the lACA or close to the axons in the mALT, and to a ventroposterior domain in the LH, along its border with the PLP (Fig. 4A-A’).

We decided to focus our tracing efforts on the lACA, a known target of the slow-cool VP3 mlALT t5ALT vPN [28, 29]. We reconstructed this PN (Fig. 2H) and discovered that 1) most of its inputs came from the simple VP3 TRN (Fig. S2B) and 2) one of its downstream targets was the uniglomerular VP2 mALT adPN (Fig. 2F). To identify the strong downstream targets of each PN (Fig. 4B), we annotated all of their presynapses and associated postsynapses in the RHS lACA (Fig. 4C), and reconstructed a random sample of these postsynaptic partners (Fig. 4D). Our sampling procedure allowed us to identify all strong downstream partners (see Methods). We also repeated this process on the LHS (Fig. S4B) and reconstructed the same types of strongly connected targets, suggesting stereotypy (Fig. S7A).

We encountered a multiglomerular VP2+ mALT adPN (Fig. 2G) downstream of both query neurons (Fig. 4E), suggesting that one function of the lACA is to permit modulation of the heating-sensitive, likely excitatory, VP2 mALT PNs by the slow cooling-responsive, likely inhibitory, VP3 t5ALT vPN. In addition, the sampled VP2 mALT adPN was presynaptic to three VP1m + VP2 mALT lvPNs (Fig. 3H-H’’).

Distinct classes of mushroom body-intrinsic neurons, or Kenyon cells (KCs), are generated sequentially during embryonic and larval development: KCγd, KCγ, KCα’/β’, KCα/βp, and KCα/β [39]. Based on light-level studies [31], we had expected to identify KCα’/β’ neurons innervating the lACA. Indeed, we reconstructed three KCα’/β’ap neurons on each side that exclusively received input from the VP3 and VP2 PNs in the lACA (Fig. 4H), strongly suggesting that there are developmentally pre-determined, dedicated thermosensory Kenyon cells. We also reconstructed one KCγd neuron on each side with a large dendritic claw in the lACA and unusually complex axon branching (Fig. 4I). This unique KCγd neuron received most dendritic inputs from the VP3 and VP2 PNs in the lACA but integrated these with additional inputs from a stereotyped lACA-associated neuron, AV3f1#2 (Fig. 5C), and from multiglomerular PNs and hygrosensory PNs in the CA (Fig. 4E) and visual PNs in the dorsal accessory calyx (dACA) (Fig. S5).

**Figure 5:**
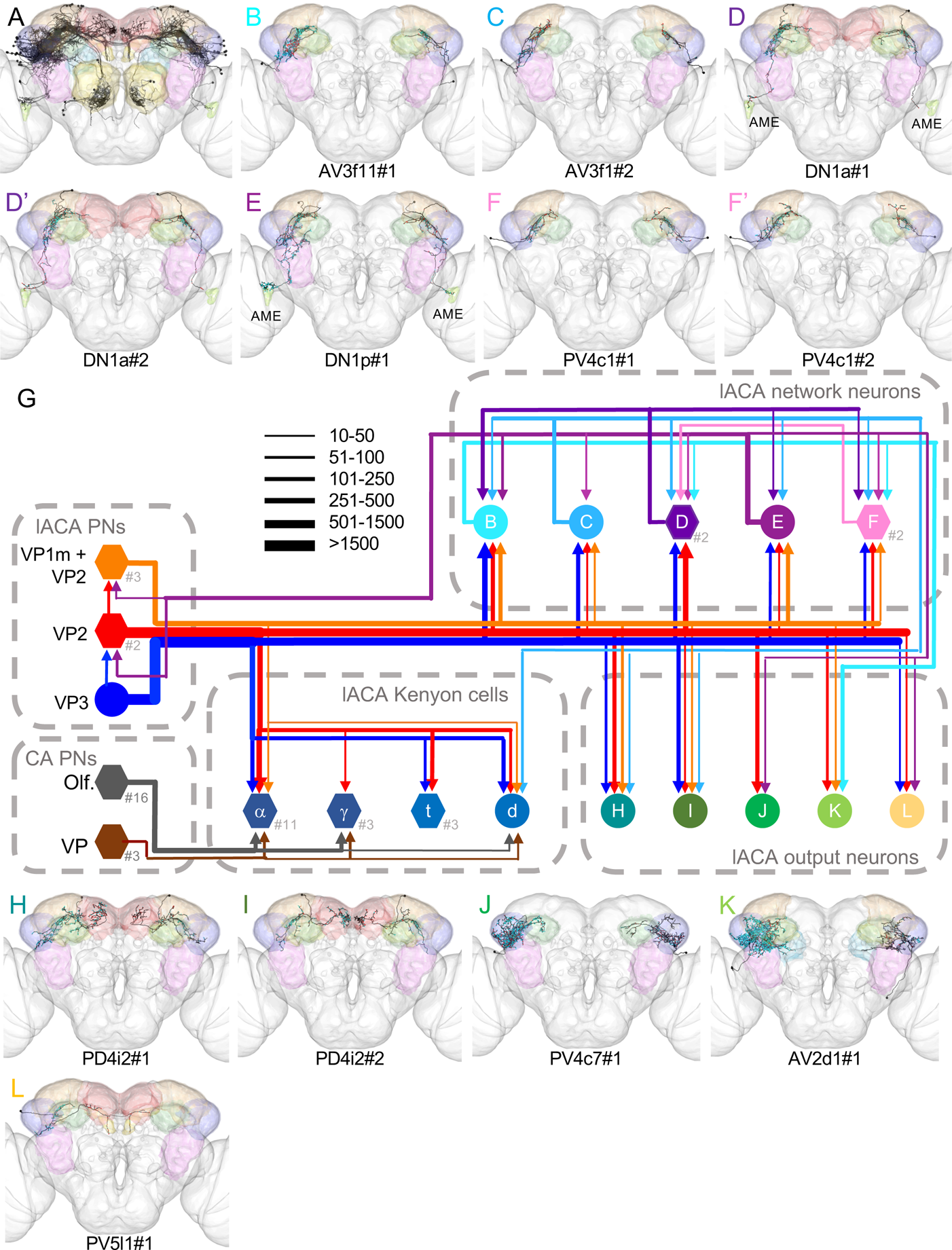
The lateral accessory calyx is a thermosensory hub with outputs to multiple neuropils. A - F. Frontal views of reconstructions of lACA target neurons. Brain neuropils have been colour-coded to correspond with all other figure panels. **A.** All targets with >10 connections to the VP3 PN and VP2 PN used for sampling. **B – F.** Strong downstream targets that form a mutually interconnected local network. **G.** lACA circuit diagram depicting connectivity between contributing VP PNs and their strong downstream targets. Letters correspond to those of panels in this figure, with lACA PNs, CA PNs, and lACA KCs carried over from Fig. 3. Circular nodes represent individual neurons, and hexagonal nodes represent pooled neurons, with the number pooled after the hashtag. Synapse numbers correlate with line thicknesses. CA PNs: Olfactory and VP PNs targeting KC dendrites in the CA. **H – L.** Frontal views of reconstructions of strong downstream targets of VP PNs in the lACA with little or no contribution to the local lACA network. For more details re: individual neurons, please see Fig. S6.

We also reconstructed 25 additional lACA-associated KCα’/β’ neurons (Fig. 4F), 24 of which belonged to the KCα’/β’ap subtype, and 15 larval-born KCγs (Fig. 4G). These two classes received additional inputs in the CA from olfactory, thermosensory, and/or hygrosensory PNs (Fig. 4E, Fig. S5). Finally, we identified one weakly connected KCγd neuron (Fig. 4J) with complex axon branching and dendritic branches in the PLP as well as in the ventral accessory calyx (vACA), a visual neuropil [40, 41]. All lACA-associated KCs originated from the same neuroblast lineage (Fig. 4K).

The main calycal inputs to lACA-associated KCs were not the same on both sides of the brain (Fig. S5). However, we calculated the expected number of PN partners of each glomerular class that would result from the random assortment of available lACA KC claws and PN boutons on the RHS (Fig. 4L) and found that lACA KCs receive disproportionately more inputs from PNs activated by odours associated with decaying fruit or danger (Fig. 4M). Inputs from VP1m + VP5 PNs (humid air) were also overrepresented (Fig. 4L). Conversely, lACA KCs received no inputs from PNs innervating pheromonal glomeruli. These results suggest that, while the synaptic partners (and thus the sensory cues) of lACA-associated KCs are not completely predetermined, they may be biased during development.

### The lateral accessory calyx is a thermosensory hub with outputs to numerous neuropils

Whilst sampling downstream of the VP3 and VP2 PNs in the lACA, we were surprised to discover that, aside from the aforementioned KCs, there were many other neurons receiving substantial input (Fig. 5A). In many cases, individual lACA target neurons (Fig. 5B, C, E, H, J, K) were unequivocally matched on both sides of the brain (Fig. S7), suggesting that they were stereotyped unique targets. In others, multiple neurons of similar morphology were recovered on both sides (Fig. 5D, D’, F, F’, I, L) but could not be individually matched (Fig. S7) without more complete knowledge of their connectivity to other up- and downstream partners.

We completely reconstructed the 12 strongest downstream targets, which received 50 - 400 combined inputs from the VP3 and VP2 PNs, and further analysed their connectivity (Fig. 5G). Many of the lACA target neurons were presynaptic to other targets (Fig. 5B - F’), forming a local lACA network, while others were only postsynaptic, likely serving primarily as output neurons to other neuropils (Fig. 5H-L). The two strongest downstream targets (Fig. 5B-C) were lACA network members and belonged to the same class of LH neurons [42], AV3f1, but could be readily distinguished based on connectivity. AV3f1#1 (Fig. 5B) was presynaptic to two classes of lACA target neurons (Fig. 5F, K) and postsynaptic to two others (Fig. D, E). AV3f1#2 (Fig. 5C) was presynaptic to AV3f1#1 and also to numerous other lACA targets (Fig. 5D, E, F, H, I), including lACA KCγdR#1 (Fig. 4I).

Three similar neurons with dorsal somas projected to the accessory medulla (Fig. 5D-E) and likely represented DN1 circadian pacemaker neurons, which can be entrained to temperature [43]. One of these (Fig. 5E) had a more posterior soma and was presynaptic to many lACA network neurons and also to its upstream lACA VP PNs. The other two (Fig. 5D-D’) were very similar in connectivity, providing input to only three classes of lACA target neurons, and likely represented the DN1a subclass [44]. The PV4c1 neurons (Fig. 5F-F’) received input from the other lACA network neurons, were presynaptic to the DN1a class (Fig. 5D), and featured arbours in the SLP.

The two lACA output neurons receiving the most input from the VP PNs were both PD4i2 neurons. One neuron projected to the border of the LH and PLP and to the SMP (Fig. 5H), a major target of MB output neurons and dopaminergic neurons [40]. The other [45] projected more medially in the SMP and featured fewer processes along the LH-PLP border (Fig. 5I). Two unique lACA output neurons had been identified in a previous study as downstream targets of DA2 PNs, an aversive olfactory channel [46]. Both ramified broadly in the LH and in the CA, although PV4c7#1 was restricted to those neuropils (Fig. 5J), while AV2d1#1 extended into the SLP, PLP, SCL, and ICL (Fig. 5K). Both received input from the VP2 PN but not the VP3 PN (Fig. 5G), and in addition, PV4c7#1 (Fig. 5J) was downstream of DN1p#1 (Fig. 5E), while AV2d1#1 (Fig. 5K) was a strong downstream target of AV3f1#1 (Fig. 5B). Finally, numerous PV5l1 neurons received input to varying degrees from the VP3 and/or VP2 PNs in the lACA. These neurons projected contralaterally, innervating the SMP and antlers of both hemispheres (Fig. 5L).

In summary, we found that six VP PNs (representing VP3, VP2, and VP1m) innervated the lACA, providing inputs to dedicated thermosensory and integrative Kenyon cells and at least seven other neuronal classes. Most lACA neurons were not connected to Kenyon cells but rather to each other, forming a local thermosensory network with outputs to circadian circuits and to brain regions implicated in innate and learned olfactory responses.

### VP PNs project to diverse targets, including at least one descending neuron

We noticed whilst sampling downstream of the VP3 and VP2 PNs in the lACA that many strong targets also received VP PN input in other neuropils. We looked downstream of the entire population of VP PNs and identified the 17 most strongly connected partners, with >150 inputs from the VP PN population. However, since reconstruction in this dataset has been focused on the LH and CA, we found that many VP PNs projecting to other neuropils shared no connections with those targets and were therefore excluded from further analysis.

Of the strongly connected population (Fig. 6A), two were themselves VP PNs taking the t2ALT, elaborating in the ventroposterior LH (Fig. 2U and Fig. 3J), and 10 received significant input in the lACA (described above, Fig. 5). We selected five of the remaining six neurons to showcase their diversity: local LH interneuron PD2c1#1 (Fig. 6B), multiglomerular projection neurons VM1+ mALT lvPN (Fig. 6C) and VP1m+ tALT mdPN (Fig. 6D), LH-SLP and SEZ connector PV9e1#1 (Fig. 6E), and descending neuron DNp44 (Fig. 6F).

**Figure 6:**
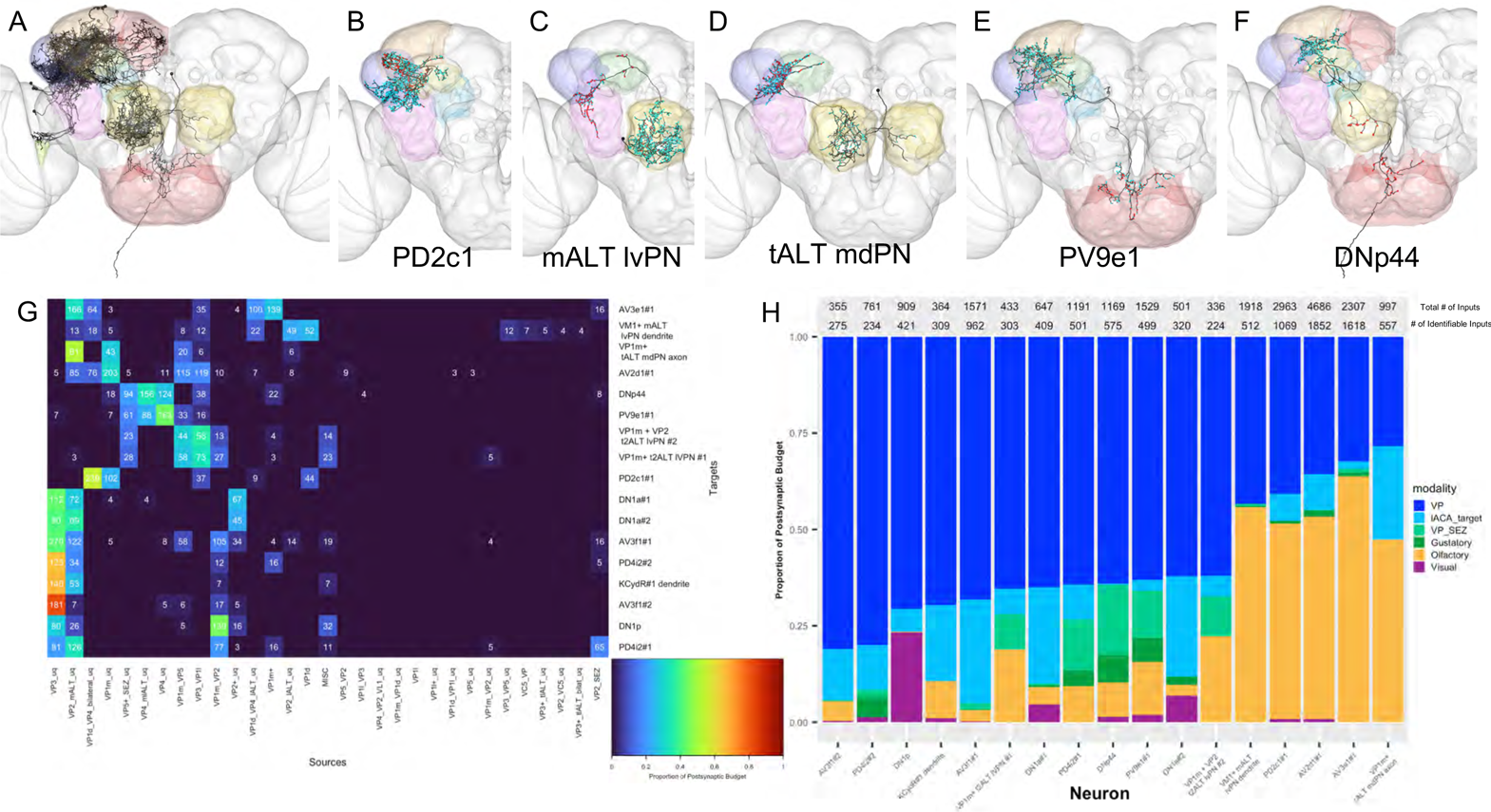
VP PNs relay information to diverse targets, including a descending neuron. A-E. Frontal views of selected strong downstream targets of RHS and bilaterally symmetric VP PNs, with brain neuropils colour-coded as in Fig. 2B’. **A.** 17 strong downstream targets of VP PNs. **B**. Local LH neuron PD2c1. **C.** Multiglomerular PN, VM1+ mALT lvPN. **D.** Multiglomerular PN, VP1m+ t-mALT mdPN. **E.** Lateral horn and SEZ interneuron PV9e1. **F.** Descending neuron DNp44. **G.** Heatmap showing contributions by VP PN classes to 17 downstream targets, normalised with total VP PN input = 1. Cell numbers denote raw connectivity numbers. Heatmap is thresholded > 2 synapses. MISC = pooled VP mPN neurons which did not show substantial connectivity. **H.** Summary stacked bar graph showing proportion of inputs from VP PNs, VP + SEZ PNs, gustatory PNs, olfactory PNs, visual PNs, and lACA target neurons, normalised with total identified inputs = 1. Total inputs and total identified inputs are shown above each bar.

We examined the specific types of VP PN input for each of these neurons (Fig. 6G) and identified some interesting patterns. The strongest lACA target neurons, including lACA KCγdR#1 (Fig. 4I), received mainly thermosensory inputs; AV3f1#2 (Fig. 5C) was almost exclusively dedicated to the slow-cool VP3 PN. While DN1a#1-2 (Fig. 5D-D’) received similar thermosensory inputs, justifying their pooling in other analyses (Fig. 5G), DN1p#1 integrated inputs from VP3 and VP1m PNs. PV9e1#1 and DNp44 innervated a similar area at the LH-PLP border (Fig. 6E-F) where both integrated input from the VP4 mlALT PNs (dry) and the VP5 + SEZ mALT PN (humid). The two VP1m+ t2ALT PNs (Fig. 2T, Fig. 3N) integrated input mainly from the VP1m + VP5 (humid) and VP3 + VP1l (cooling) bilateral mALT PNs. The VP1m+ tALT mdPN received input from the VP2 mALT adPN (heating) and the VP1m lALT lPN, while the VM1+ mALT lvPN received input mainly from the VP2 lALT PN (heating). Finally, AV2d1#1, and PV4c7#1 were mixed, integrating VP2 (heating) with VP1m (suspected humid) and the bilateral VP1d and VP4 PN (evaporative cooling/dry).

Next, we characterised the input to each of these top 17 targets by modality. We focussed on identifiable inputs and separated them into VP PNs, VP PNs with SEZ innervation, 3rd order lACA target neurons, olfactory, gustatory and visual categories. VP PNs provided between 28% and 81% of the known input to these 17 targets, all of which integrate information from multiple modalities (Figure 6H). The extent of cross-modal integration varied by neuron; for example, the PD2c1#1 local interneuron, the VM1+ mALT lvPN dendrite, and the VP1m+ tALT mdPN axon all integrated VP and olfactory information, with negligible input contribution from gustation. In contrast, PV9e1#1 and DNp44 integrated information from all 6 categories, with thermo/hygro input from VP PNs and lACA target neurons occupying >50% of identifiable input. Lastly, DN1p and DN1a neurons receive a substantial proportion of visual input, connecting medulla projection neurons via the accessory medulla to the lACA, thereby linking thermoreception with photoreception in the circadian clock.

## DISCUSSION

Neural circuits that detect and respond to temperature and humidity are vital for survival and reproduction, but they are much less well-characterised than the olfactory system, with which they share many anatomical correlates in insects. In *Drosophila*, many thermo- and hygrosensory neurons are housed in the antennae, innervate dedicated glomeruli in the antennal lobe, and target stereotyped projection neurons that relay information to higher brain centres [47]. By reconstructing individual neurons and their synapses in a whole brain EM volume, we have identified novel primary, secondary, and tertiary neurons.

We reconstructed sensory neurons in seven VP glomeruli, two of them novel; surprisingly, the average number per VP glomerulus was only about 1/4 that of olfactory glomeruli [48]. Based on their ionotropic receptor expression, we suggest that the novel glomeruli represent cool/cooling (VP1l) and humidity (VP1m), but confirmation awaits future physiological and behavioural experiments. Our VP1d most likely corresponds to the glomerulus originally designated VP1, which responds weakly to cooling, dry, and ammonia and may represent evaporative cooling.

Projection neurons relay information from the antennal lobe glomeruli to higher brain centres, where they connect to partners critical to innate and learned behaviour. We reconstructed 19 novel VP PN types using our systematic connectomics approach; further studies with sparse driver lines will be necessary to define their valences and functions. Unexpectedly, and unlike olfactory PNs, many VP PNs received substantial input from two distinct glomeruli. In most cases, both were VP glomeruli, but VC5 was innervated by several PNs in this study, including two resembling a reported thermosensory neuron type [30], hinting that this medial, ventroposterior glomerulus might also be thermosensory. Biglomerular PNs typically co-innervated VP glomeruli of similar, rather than opposing, valences, suggesting a pooling of related sensory information. Integration of antagonistic inputs (e.g. heating + cooling) or across modalities was more characteristic of third order neurons.

By characterising the connectome of the lACA, we identified three α’/β’ Kenyon cells specialised for thermosensation as well as many others that integrated thermosensory with olfactory and/or hygrosensory information. Future analysis of downstream partners should reveal whether either of these classes connects to specific subsets of dopaminergic neurons (DANs) and/or mushroom body output neurons (MBONs) to influence memory formation or retrieval. The two lACA-associated KCγd neurons are distinct from other KCγds, which project to the γ lobe dorsal tip [37], in that they exhibit highly branched axons wrapping around the γ lobe, suggesting that they output onto distinct synaptic partners.

We also identified 8 new classes of lACA-associated neurons, most of which connect, not to Kenyon cells, but rather to each other and/or as yet unidentified neurons in other neuropils. Intriguingly, we discovered circadian clock neurons DN1a and DN1p innervating the lACA, thus potentially mediating temperature-based entrainment [49] or, alternatively, modulating temperature preference, which fluctuates with a daily rhythm in animals from flies to humans [50]. Three of the lACA network’s main output neuropils are the LH, SMP, and antlers, all previously associated with olfactory responses [51]; in particular, the SMP features both DAN inputs and MBON outputs [40]. However, the specific synaptic partners of lACA output neurons have yet to be identified.

We find that while thermo- and hygrosensory inputs are initially segregated by valence in the antennal lobe, they are integrated with each other and with olfactory inputs at higher levels. While we can only speculate regarding the importance of this integration, it seems reasonable to hypothesise that the rate of encountering odour molecules changes with varying temperature and humidity, requiring some knowledge of context for accurate odour perception. It is also plausible that antennal thermosensation and hygrosensation are more ancient, with a system for detecting, remembering, and responding to specific volatile chemicals only later being layered on top.

Whilst this study has significantly improved our picture of thermo- and hygrosensory circuitry, it is clear that many gaps remain at the level of third order target neurons in higher brain areas. Given the time-consuming labour of manual reconstruction, our comprehensive efforts at this level were restricted to the lACA. Taking full advantage of whole EM volumes to elucidate the organisation of brainwide circuits, and making comparisons between animals to assess natural and experimental variation, will require rapid, accurate automated segmentation of neuronal profiles and their synapses [52]. This study should provide a lasting platform for this next stage of exploration and the functional studies that will accompany them.

## ACKNOWLEDGMENTS

This work was supported by a Wellcome Trust Collaborative Award (203261/Z/16/Z to G.S.X.E.J, M.L. and D.B.), a National Institute of General Medical Sciences training grant (T32 GM07122, T.S.), grants from the National Institute of Allergy and Infectious Diseases (R01 AI122802 and R21 AI40018) and National Science Foundation (IOS 1557781) to P.A.G.; MRC LMB Graduate Studentship to M.W.P.; a Boehringer Ingelheim Fonds PhD Fellowship and a Herchel Smith Studentship to A.S.B, a Cambridge Neuroscience-PSL collaborative grant supported by the Embassy of France in London to G.S.X.E.J. and core support from the MRC (MC-U105188491) to G.S.X.E.J.

We thank Zachary Knecht for preliminary experiments and assistance with identification of thermo- and hygrosensory glomeruli and Shigehiro Namiki for assistance with identification of DNp44. We thank all CATMAID tracers from the FAFB community who contributed to the neuron reconstructions featured in this study, especially Imaan Tamini, Ilenia Salaris, Nik Drummond, Lucia Kmecova, and Jawaid Ali. We also thank all members of the Jefferis and *Drosophila* Connectomics groups who shared R or Python code that was modified for use in this study, particularly Kimberly Meechan, Fiona Love, Amelia Edmondson-Stait, and Istvan Taisz.

## DECLARATION OF INTERESTS

The authors declare no competing interests.

## METHODS

### EM tracing

Neuron skeletons were traced in a full adult female *Drosophila* brain EM volume either manually as previously reported [32], or by automated segmentation, using a modified version of CATMAID [53, 54]. Skeleton fragments that had been automatically segmented with the flood-filling technique [52] were manually joined in a separate CATMAID instance, imported into the manual CATMAID instance, and merged with manually traced partial skeletons where possible (https://github.com/flyconnectome/fafbseg-py). Skeletons were either traced to identification (with soma, backbone, and main branches) or to completion (with all twigs, including pre- and postsynapses, and fully reviewed [36].

Synapses were annotated based on previously described criteria: thick, dark active zone, presynaptic membrane specialisations (T-bars, vesicles), and a synaptic cleft [55]. We scored each continuous synaptic cleft as a single presynapse and opposing neuronal membranes in contact with that synaptic cleft as associated postsynapses.

### Analysis and representation of traced skeletons

The connectivity and graph widgets in CATMAID were used to analyse and visualise connectivity between traced skeletons. R packages from the natverse collection (http://natverse.org/) [56] were used to plot traced skeletons and analyse their morphology. Custom R and Python scripts were used to perform additional analysis, as described below.

### Neuron clustering via NBLAST

The nat.nblast R package (https://github.com/natverse/nat.nblast) was used to compare neuron skeletons by morphology and position and generate a hierarchical clustering of the neurons within each group (e.g., VP PNs), providing a method for identifying neuron types [33].

### Light vs EM comparisons of neuron morphology

To assist in the identification of our reconstructed neurons, we used NBLAST to compare them with segmentations of annotated PNs in the light-level FlyCircuit database [33, 57] (http://www.flycircuit.tw/). We used linear and then non-rigid transformations to bring neurons from FAFB into FCWB (Fly Circuit whole brain) space, then performed an all-by-all NBLAST to look for the closest matches. Neurons that did not have clear matches in FlyCircuit could sometimes be manually identified by comparing morphology to published data. Sparse driver lines predicted to label our reconstructed neurons were identified in a similar way through comparisons with the Fly Light database (http://flweb.janelia.org/cgi-bin/flew.cgi).

### Neuronal nomenclature

Sensory neurons were named for the AL glomerulus they innervated. Projection neurons were named according to their AL innervation, AL tract, and neuroblast lineage [35]. Kenyon cells were named according to their positions in the peduncle and the MB axon lobes they innervated [40]. Lateral horn neurons were named for their primary neurite tracts and areas of innervation, according to previously established conventions [42].

### EM downstream sampling

Postsynapses downstream of the VP3 mlALT t5ALT vPN and VP2 mALT adPN in the lACA were manually annotated, then randomly sampled and reconstructed to identification. We continued sampling until the 10 most recent novel hits featured fewer than five connections with the query neuron (Fig. S4a), corresponding to 21 - 30% of postsynapses for each PN. All identified neurons and orphan fragments were rechecked, and the 12 most strongly connected downstream targets were traced to completion and reviewed.

### Analysis of Kenyon cell inputs

The dendrites of lACA-associated KCs on the RHS of FAFB were completed in CATMAID, and their postsynapses were manually associated with upstream neurons - mainly RHS or bilaterally symmetric antennal lobe PNs, which had already been traced to identification in this dataset [32, 36]. Analysis of KC inputs was performed in R with a customised script, utilising the nat and elmr packages from the natverse. KC skeletons were bridged from FAFB space into the JFRC2 template [28] (http://www.virtualflybrain.org/ using the <u>nat.flybrains</u> package and then pruned with the split_neuron_local() function in the <u>tracerutils</u> package to include only dendrites (branches off the main tract in the calyx). For each KC branch, all upstream PNs contributing at least 5 synapses were identified and then weighted equally as inputs observed, independent of synapse count or bouton size. (Non-PN partners and the VP3 and VP2 PNs originally used for sampling were omitted from further analysis.)

Expected numbers of inputs representing each AL glomerulus were based on the total number of boutons reported [32], or newly identified, for each PN class. PN boutons were manually annotated and extracted using the rmushroom package (https://github.com/zhihaozheng). A virtual KC population with the same number of KCs and PN inputs was created by random draw, with the probability of each input based on the percentage share of associated PN boutons. This was repeated 1000 times to generate an average number of expected inputs for the population. Inputs from each glomerulus were assigned a putative valence based on the olfactory literature [36].

### *Drosophila* strains and driver line generation

Ir40a-LexA[27], Gr28b.d-Gal4 [58], UAS-myr:GFP (P[10UAS-IVS-myr::GFP]attP1) [59] were previously described. LexAop-RFP (P[lexA-2xmRFP.nls]2) (RRID:BDSC 29956) was obtained from Bloomington Stock Center.

Ir21a-T2A-Gal4 and Ir68a-T2A-Gal4 where generated using techniques previously described [60]. Briefly, intron regions in the Ir21a and Ir68a genes were targeted via homology directed repair by injecting two plasmids, one containing gRNA (Ir21a: 5’-GCGCGTGAGTATTGCTTAAT; Ir68a: 5’-GTGTGATATGAAAAGCATTC) another with T2A-Gal4 3XP3-RFP at phase0 (flanked by homology arms; Ir21a: 5’arm primers 1-2, 3’arm primers 3-4; Ir68a: 5’arm: primers 5-6, 3’arm primers 7-8) into embryos carrying Vasa-Cas9 3XP3-EGFP (kindly shared by Kate Koles and Rodal Lab,-----Brandeis University).

#### Primers used

Primer 1: 5’-atacgactcactatagggcgacccaccggtCGATATGTCATATTATTGGGTAGCTCTGGT

Primer 2: 5’-ccgaaaaccgcttctgacctggggcggccgcAAGCAATACTCACGCGCAT

Primer 3: 5’-ccgaaaaccgcttctgacctgggggcgcgccAATAGGGATACGTTTTTGTAACAAATAATGCGC TTCACACAGGAG

Primer 4: 5’-aagggaacctccccactagtggtaccCTCAAATGAAGCGCCGATCGG

Primer 5: 5’-atacgactcactatagggcgacccaccggtGGAACCTCTTTGCCAGTTGCC

Primer 6: 5’-ccgaaaaccgcttctgacctggggcggccgcTGCTTTTCATATCACACTAGATTATTTTG

Primer 7: 5’-ccgaaaaccgcttctgacctgggggcgcgccTTCCGGCGACTTAATGGCTTTGT

Primer 8: 5’-aagggaacctccccactagtggtaccCTTCTAAAGAGATGGCCAAGCAAAAGC

### Immunohistochemistry

For brain dissections, whole flies were fixed for 2hrs in 4% paraformaldehyde while rotating and washed several times prior to dissections. The dissected brains were incubated at room temperature in blocking solution (10% normal goat serum) for one hour. Primary and secondary antibody incubations were done at 4°C for 48hrs each. Washing between antibody incubations was performed over 24hrs at 4°C.

Third antennal sections were dissected, fixed in 4% paraformaldehyde for 1hr on ice, and incubated in blocking solution for 1hr at room temperature. Primary and secondary antibody incubations were done at 4°C for 24hrs each while rotating. Washing between antibody incubations were performed at room temperature 3-4 times with PBS-Tx (PBS with 0.5% Triton X).

The following antibodies were used: mouse anti-nc82 at 1:200 (Developmental Studies Hybridoma Bank), mouse anti-Elav at 1:30 (Developmental Studies Hybridoma Bank), chicken anti-GFP at 1:100 (Avēs Labs), rabbit anti-DsRED at 1:200 (Takara Bio), goat anti-mouse Cy3 at 1:200 (Jackson Labs), goat anti-mouse Cy5 at 1:200 (Jackson Labs), goat anti-chicken 488 at 1:200 (Life Technologies), goat anti-rabbit Cy3 at 1:200 (Jackson Labs).

### Confocal microscopy

Imaging of antennae and the antennal lobe were performed using a Zeiss LSM 880 confocal microscope with a 63X oil lens. Maximum intensity z-stack projections were made from z-stack sections taken every 1µm.

---

**Figure S1.**
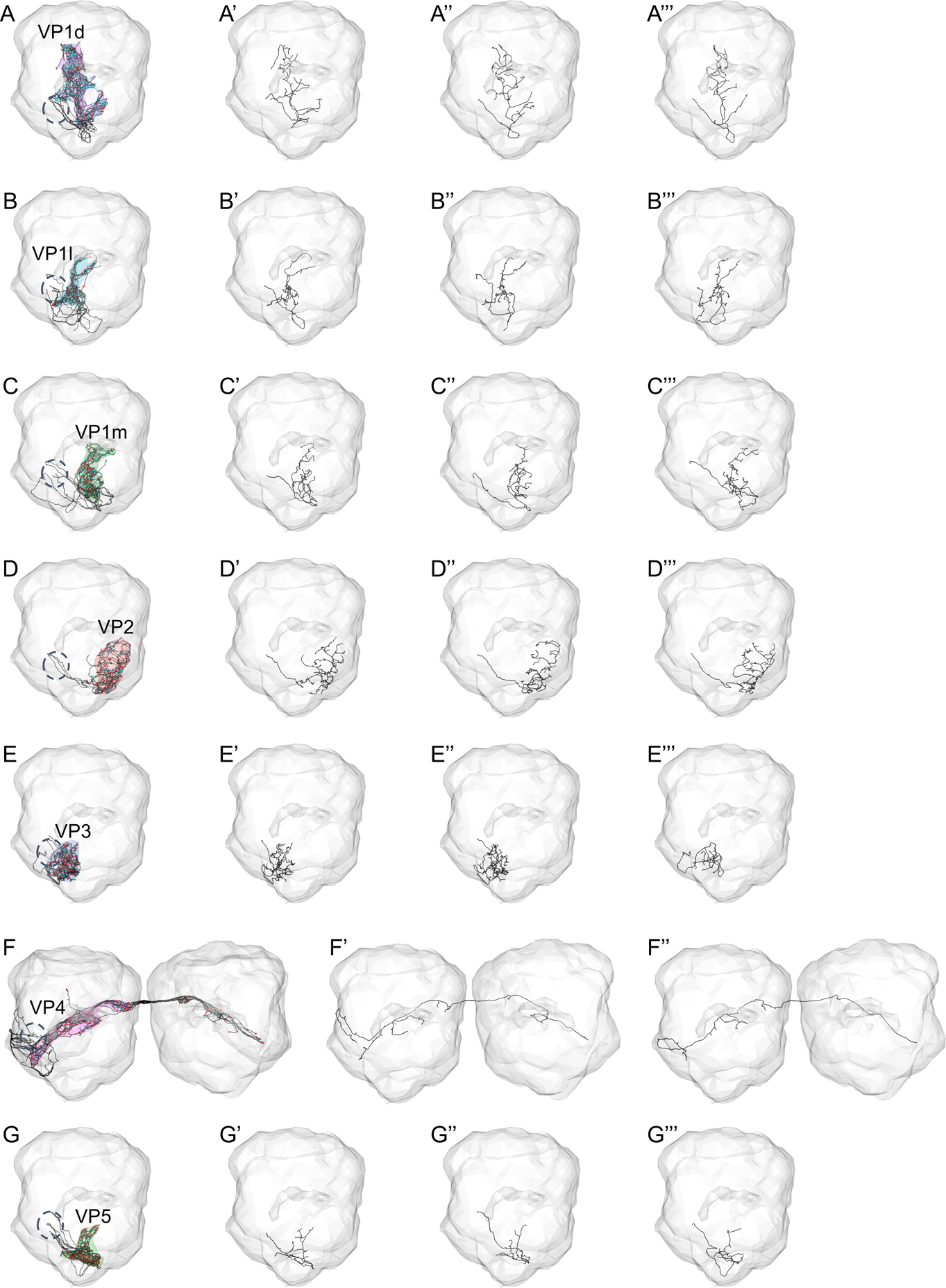
Individual reconstructions reveal morphological stereotypy for seven distinct classes of VP sensory neurons. Right antennal lobe, frontal view. Reconstructed neurons in black, with presynaptic sites in red and postsynaptic sites in cyan. Dashed circle: right antennal nerve. **A – A’’’.** Full population (9) and three completed examples of RHS VP1d HRNs. **B – B’’’.** Full population (6) and three completed examples of RHS VP1l HRNs. **C – C’’’.** Full population (6) and three completed examples of RHS VP1m HRNs. **D – D’’’.** Full population (3) and three completed examples of RHS VP2 TRNs. **E – E’’’.** Full population (4) and three completed examples of RHS VP3 TRNs. **E’’’** depicts the simpler, suspected non-aristal, TRN. **F – F’’’.** Full population (14) and two completed examples of RHS VP4 HRNs. **G – G’’’.** Full population (9) and three completed examples of RHS VP5 HRNs.

**Figure S2.**
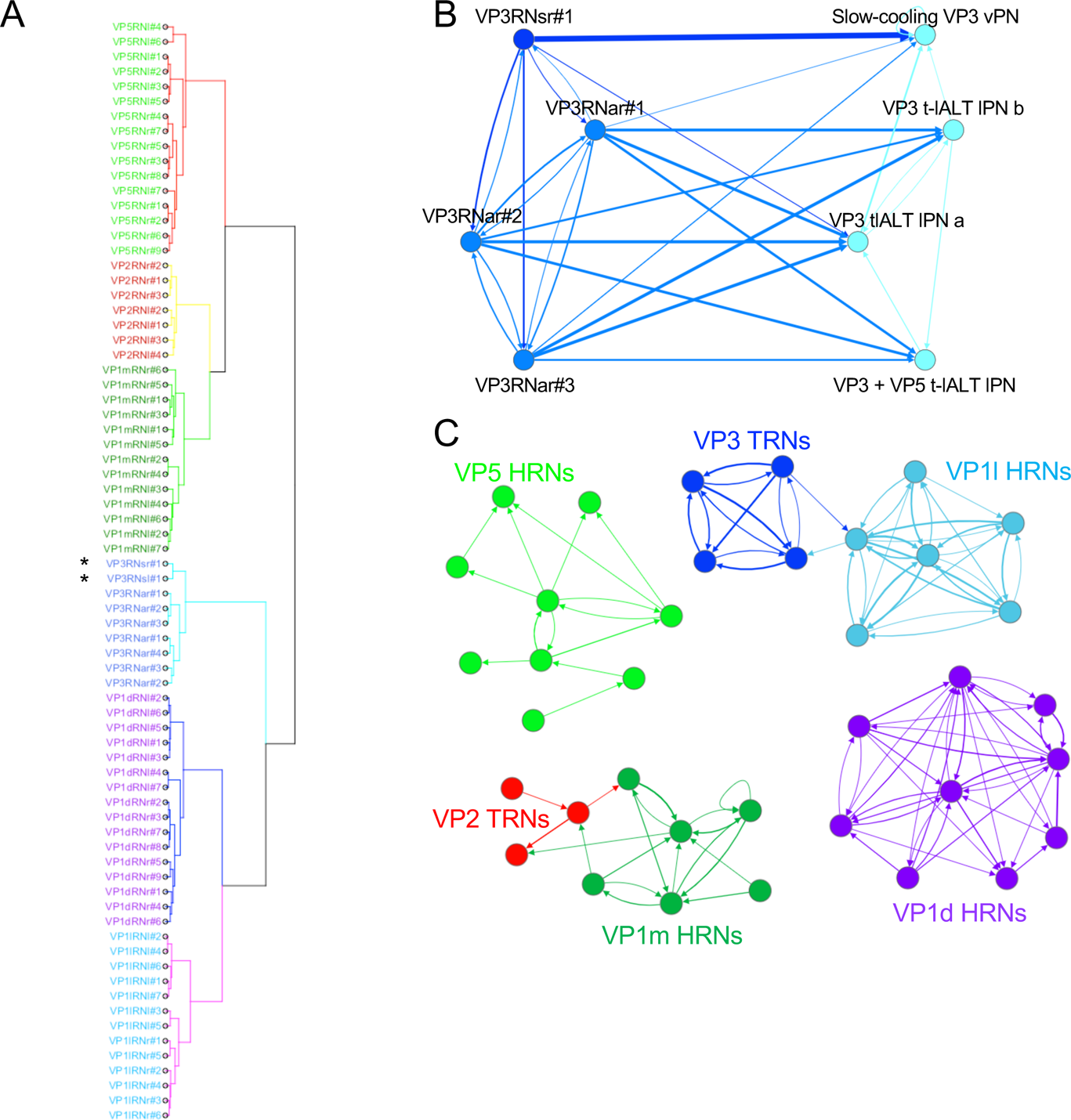
There are seven distinct populations of thermosensory and hygrosensory neurons. **A.** Dendrogram clustering the unilateral sensory neurons in each hemisphere by morphology and position. Neuron names are colour-coded by glomerulus. Asterisks: simple VP3 TRNs. **B – C.** Connectivity graphs for thermosensory and hygrosensory neuron populations. Arrow directions and thicknesses reflect direction of information flow and numbers of synaptic connections. Each circle (node) represents one neuron, colour-coded by class. **B.** The unusually simple VP3 TRN provides input to the slow-cool VP3 PN but not fast-cool VP3 PNs. Dark blue: simple VP3 TRN and slow-cool VP3 PN. Medium blue: remaining three (putative aristal) VP3 TRNs. Light blue: VP3 PNs taking the t-lALT. Edges of <3 connections have been omitted for visual clarity. **C.** VP1d, VP1l, and VP1m sensory neurons constitute three unconnected populations.

**Figure S3.**
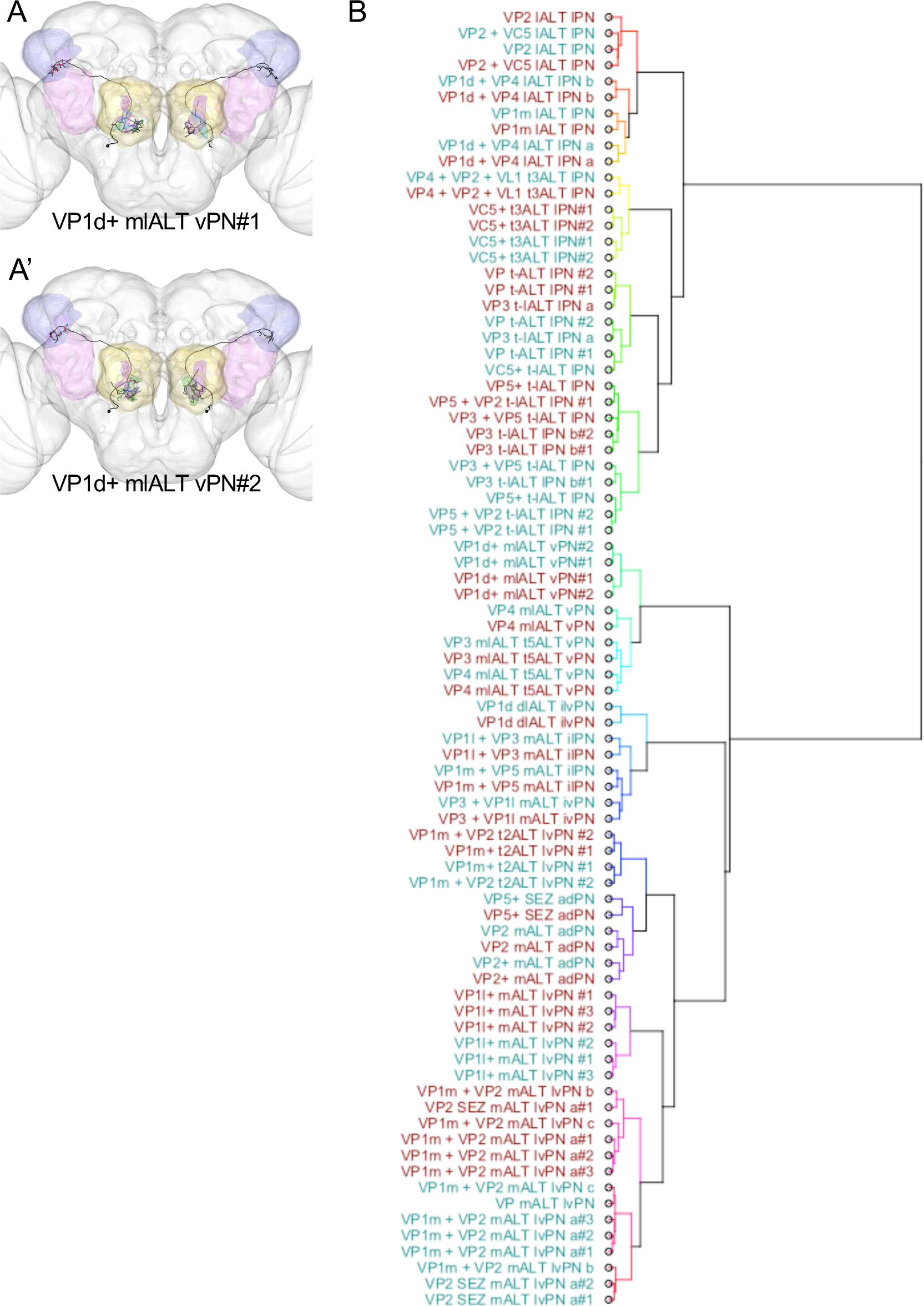
Individual VP PNs and classes of VP PNs are similar on the RHS and LHS, supporting bilateral stereotypy. A – A’. Frontal views of VP1d+ mlALT vPN reconstructions in FAFB, to be excluded from further analysis. Brain neuropils have been colour-coded to correspond with Fig. 2B’ and all other figure panels. **B.** Dendrogram comparing VP PNs in each hemisphere by position and morphology. Dark cyan: RHS. Dark red: LHS.

**Figure S4.**
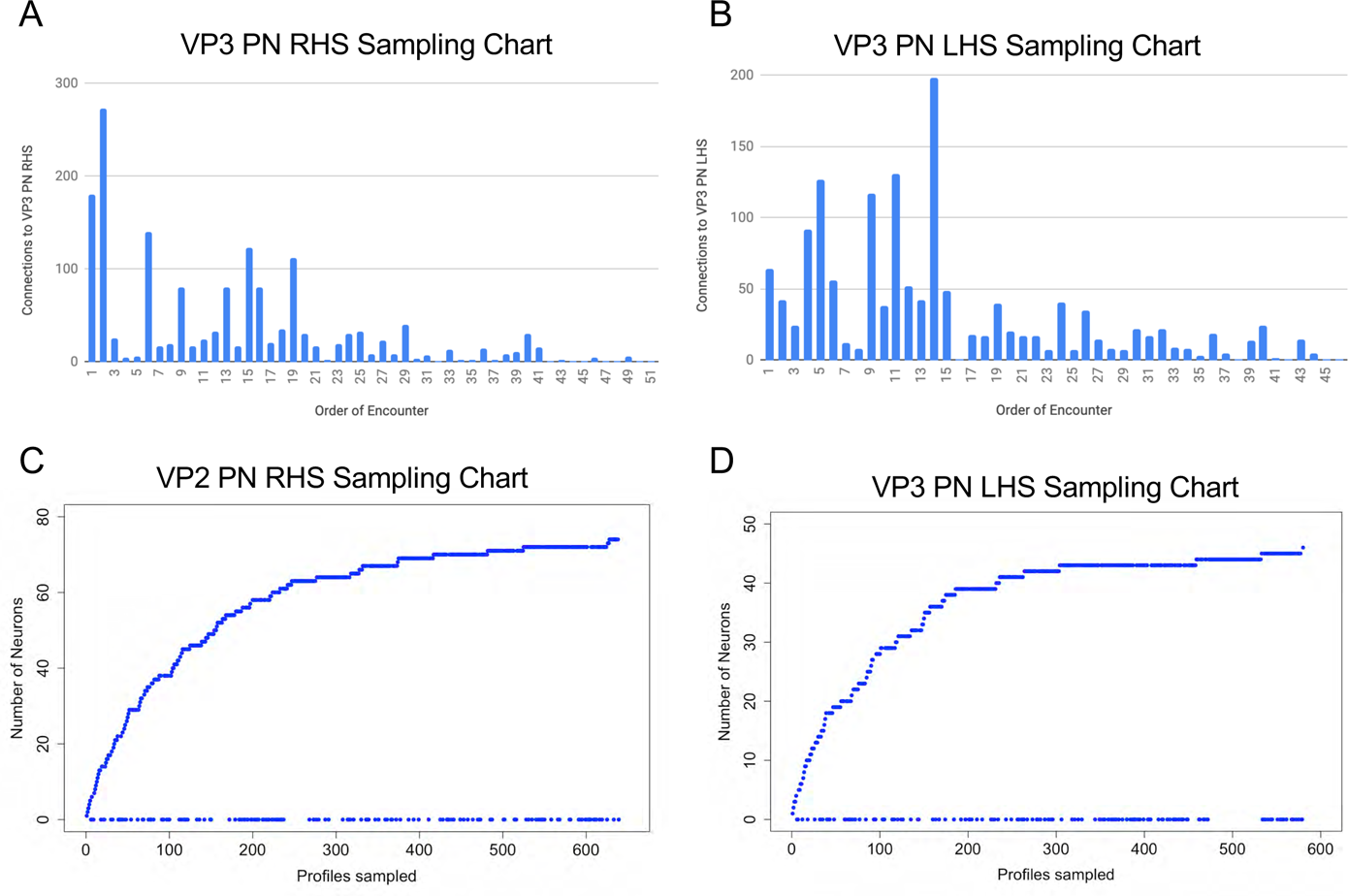
Random sampling from the VP3 mlALT t5ALT and VP2 mALT PNs identified all strongly connected downstream partners. A - B. Histogram depicting connection strength of recovered novel hits from successive sampling of randomised postsynaptic profiles downstream of **A.** the RHS VP3 mlALT t5ALT PN and **B.** the LHS VP3 mlALT t5ALT PN. **C - D.** Sampling curves depicting total number of unique neurons recovered from successive sampling of randomised postsynaptic profiles for **C.** the RHS VP2 mALT PN and **D.** the LHS VP3 mlALT t5ALT PN.

**Figure S5.**
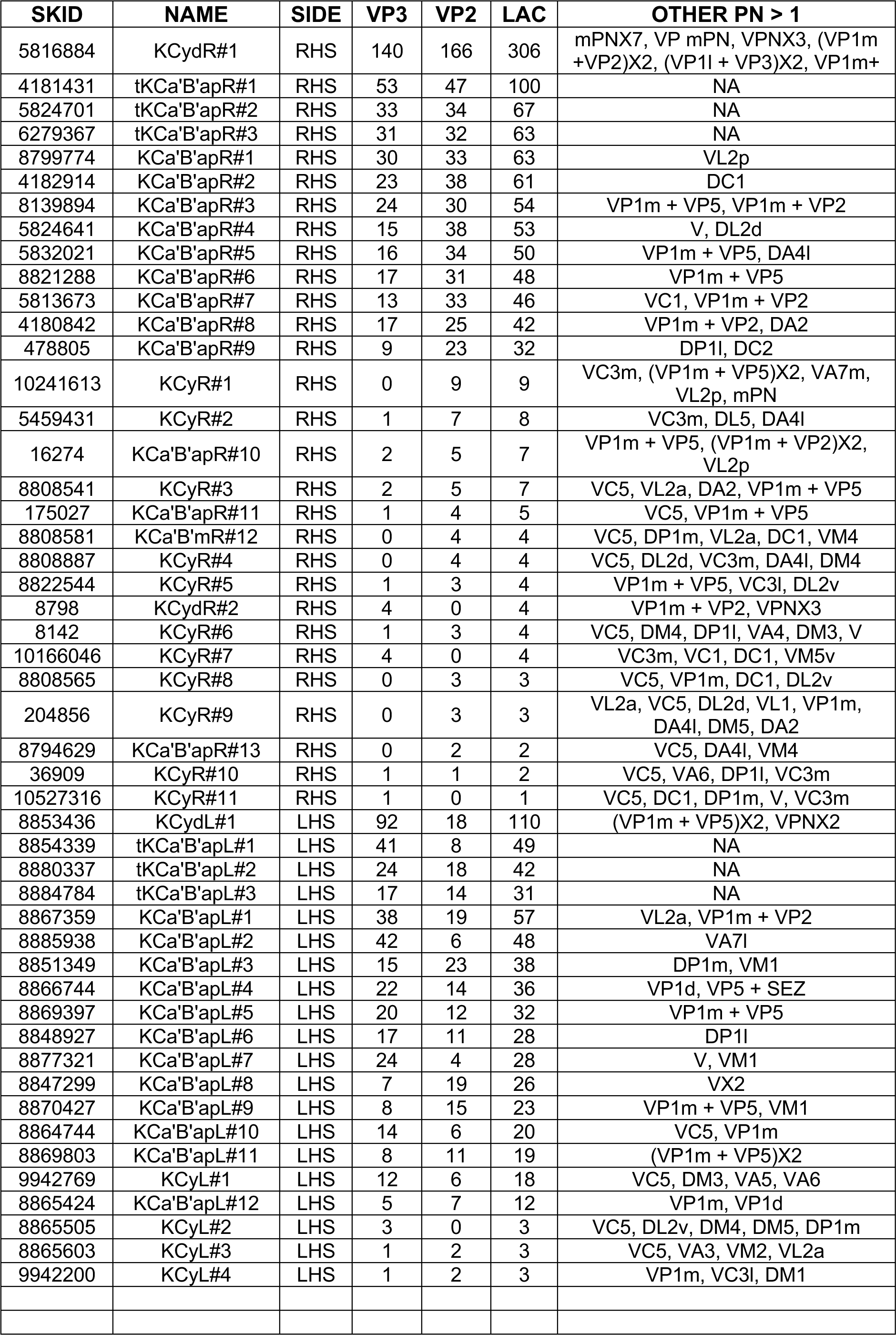
Table of lACA-associated Kenyon cells and their inputs SKID: skeleton ID. NAME: our designation, based on Kenyon cell class and hemisphere. SIDE: hemisphere. VP3, VP2, and LAC: Numbers of inputs from the VP3 PN, VP2 PN, and both PNs, respectively. OTHER PN >1: other projection neurons providing input, labeled by class. mPN: multiglomerular PN. VPN: visual PN.

**Figure S6.**
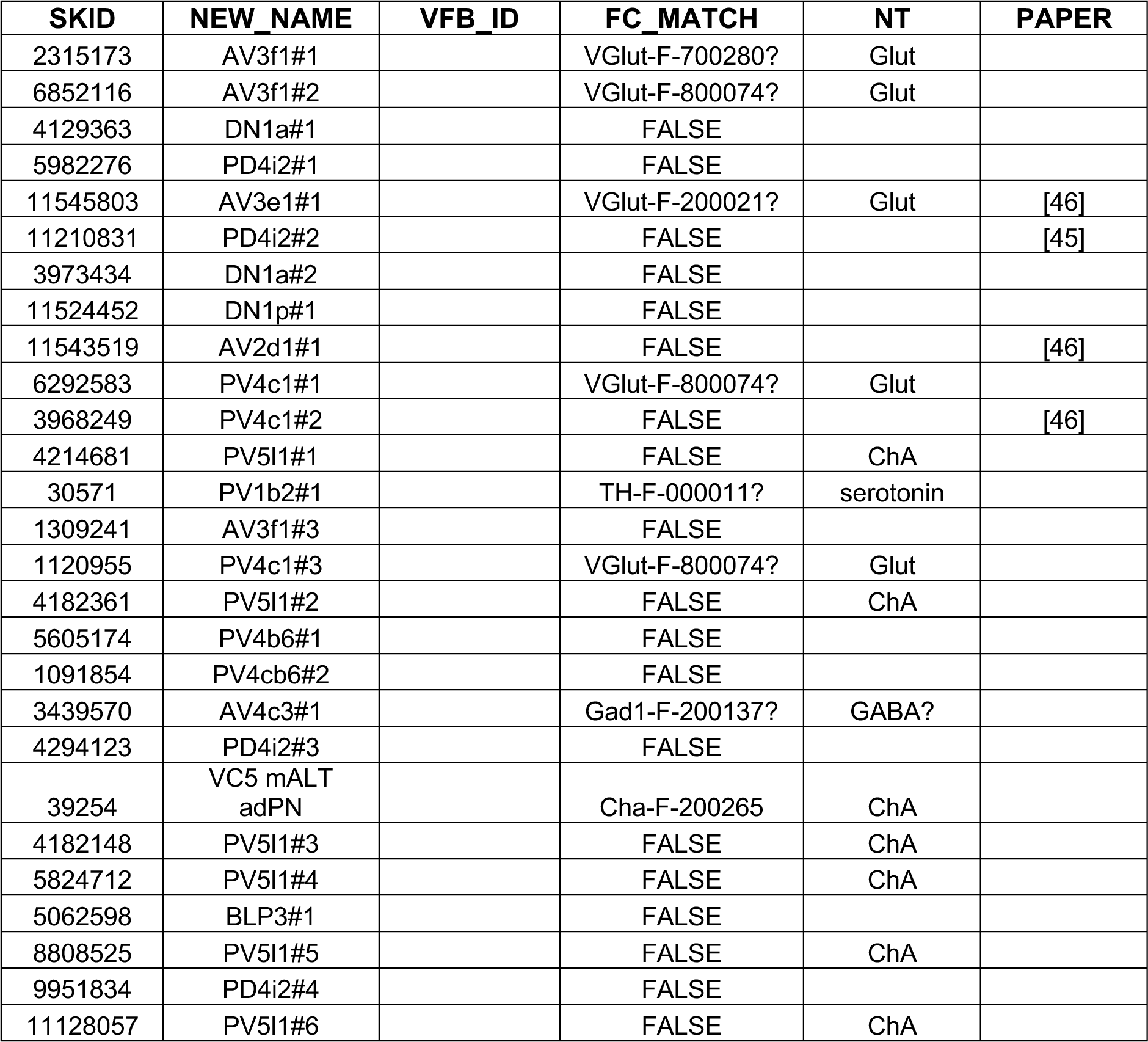
Table of non-Kenyon cell lACA neurons SKID: skeleton ID. NEW_NAME: our designation, based on neuroblast lineage (Ito) or lateral horn tract (Frechter). VFB_ID: VFB identification number. FC_MATCH: corresponding single-cell clone in FlyCircuit, based on NBLAST. NT: putative neurotransmitter, based on FlyCircuit driver. PAPER: Previous publications featuring reconstructions of this neuron in FAFB.

**Figure S7.**
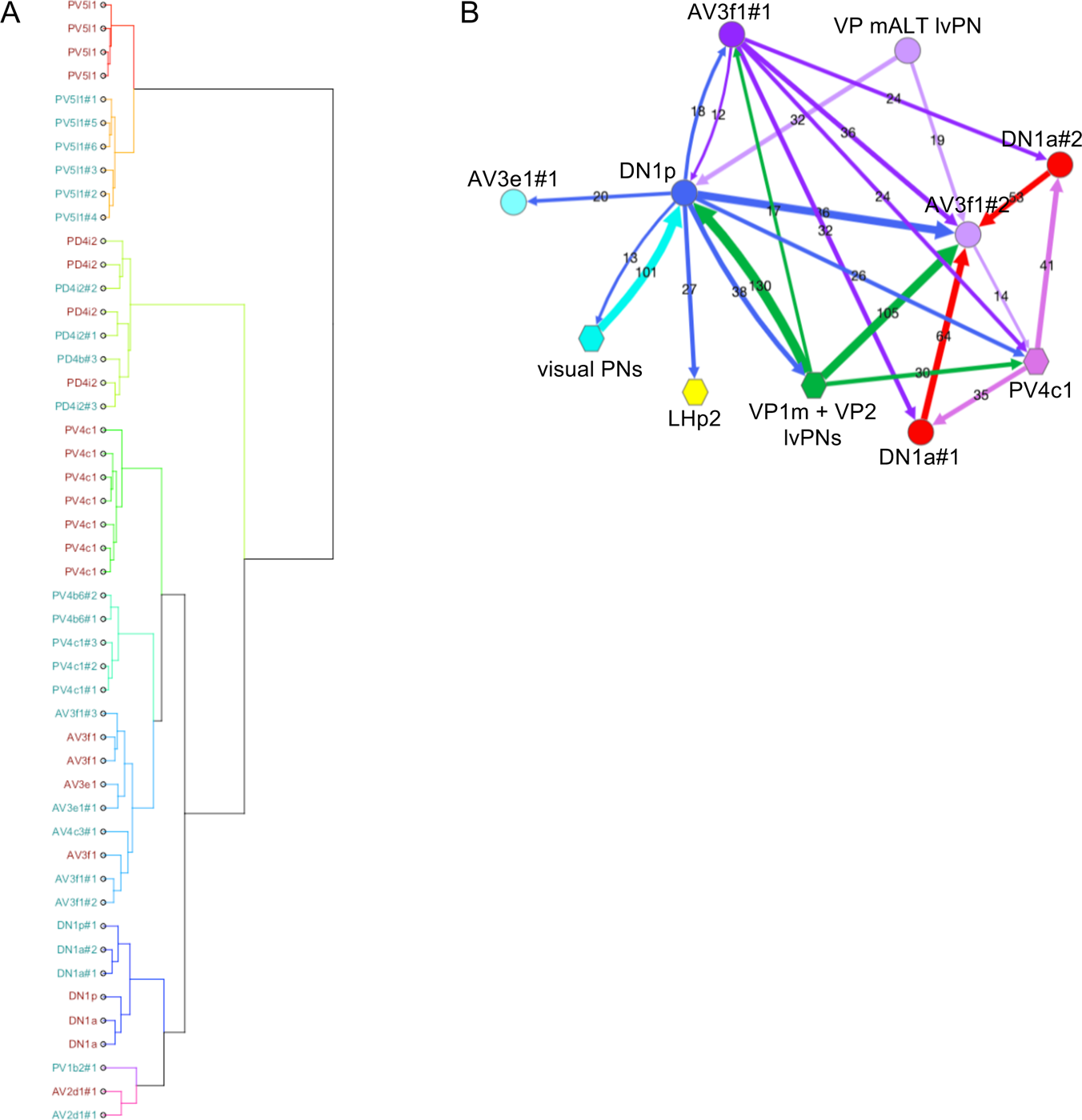
Non-Kenyon cell lACA neurons. **A.** Dendrogram clustering the targets of the VP3 and VP2 PNs in the lACA by position and morphology using NBLAST. Dark cyan: RHS targets. Dark red: LHS targets. **B.** Connectivity graph showing that DN1a#1 and DN1a#2 have similar connectivity, distinct from that of DN1p. Each circle (node) represents one neuron and each hexagon represents multiple pooled neurons of the same class. Arrow directions and thicknesses reflect direction of information flow and numbers of synaptic connections. Edges of <10 connections and the VP3 and VP2 PNs used for sampling have been omitted for clarity.

## Notes

#### Summary of Updates

Single pdf; minor fixes.

